# Identification of drug-like molecules targeting the ATPase activity of dynamin-like EHD4

**DOI:** 10.1101/2024.04.11.589129

**Authors:** Saif Mohd, Andreas Oder, Edgar Specker, Martin Neuenschwander, Jens Peter Von Kries, Oliver Daumke

## Abstract

Eps15 (epidermal growth factor receptor pathway substrate 15) homology domain-containing proteins (EHDs) comprise a family of eukaryotic dynamin-related ATPases that participate in various endocytic membrane trafficking pathways. Dysregulation of EHDs function has been implicated in various diseases, including cancer. The lack of small molecule inhibitors which acutely target individual EHD members has hampered progress in dissecting their detailed cellular membrane trafficking pathways and their function during disease. Here, we established a Malachite green-based assay compatible with high throughput screening to monitor the liposome-stimulated ATPase of EHD4. In this way, we identified a drug-like molecule that inhibited EHD4’s liposome-stimulated ATPase activity. Structure activity relationship (SAR) studies indicated sites of preferred substitutions for more potent inhibitor synthesis. Moreover, the assay optimization in this work can be applied to other dynamin family members showing a weak and liposome-dependent nucleotide hydrolysis activity.

## Introduction

Dynamin superfamily proteins are involved in various cellular membrane remodeling events [1]. They harness energy from nucleotide hydrolysis to perform mechano-chemical work. The proteins oligomerize on the surface of membranes, leading to stimulation of their low basal nucleotide hydrolysis activity [2]. While most dynamin superfamily proteins are GTP-binding and hydrolyzing enzymes, Eps15-homolgy domain-containing proteins (EHDs) comprise a highly conserved dynamin-related ATPase family, with four members in eukaryotes (EHD1-4) [3]. As other dynamin superfamily members, EHDs bind to liposomes and oligomerize in ring-like structures at their surface, inducing liposome tubulation [4–9]. EHD oligomerization leads to an ∼10-fold stimulation of the low intrinsic ATPase activity [4, 7, 9].

EHDs are implicated in a variety of cellular membrane trafficking pathways. For example, EHD1 and EHD3 localize to the endocytic recycling compartment and mediate the release of cargo molecules en route to the cell surface [10–13]. The proteins also play a role in ciliary membrane trafficking and sonic hedgehog signaling [14, 15]. EHD2 localizes to caveolae, which are bulb-shaped invaginations of the plasma membrane [5, 16, 17]. EHD2 is thought to assemble into a ring-like oligomer at the neck of caveolae, thereby stabilizing them to the plasma membrane [18, 19]. EHD4, formerly known as Pincher, mediates retrograde endosomal trafficking of neurotrophin receptors [20, 21] and also participates in primary ciliogenesis [22].

Dysregulation of EHDs’ function has also been implicated in human disease (reviewed in [3]). For example, EHD1 is overexpressed in several cancers, which appears to mediate chemotherapy resistance, epithelial-mesenchymal transition and stem cell behavior [23–25]. Mechanistically, EHD1 promotes the trafficking of insulin-like growth factor 1 receptor and other receptor tyrosine kinases to the cell surface, therefore promoting tumorigenesis and metastasis [26]. Mutations in EHD1 are associated with tubular proteinuria and deafness in human [27]. EHD2 knock out mice have increased fat depositions around organs and obese individuals show significantly lowered EHD2 expression levels [19]. Furthermore, overexpression of EHD2 was shown to promote tumorigenesis and metastasis in triple negative breast cancer [28]. Strongly decreased EHD3 expression levels were found in oral squamous cell carcinoma [29] and gliomas [30], while EHD3 overexpression was found in a rat model of ischemic heart failure [22]. EHD4 was shown to regulate urinary water homeostasis [31]. Given the involvement of EHDs in various physiological processes and its dysfunction in pathologies, small molecules that regulate EHDs’ activity could be useful tools to study their function in disease and potentially ameliorate some of the disease phenotypes.

Previously, EHD ATPase activity was determined via a High Pressure Liquid Chromatography (HPLC)-based method [4, 7]. However, the HPLC setup and assay parameters are not suitable for high throughput drug screening (HTS). Another study used a Malachite green-based assay for EHD1 and EHD2 [9], but at relatively high protein and substrate concentrations, which prevents direct application in HTS. Various dynamin drug screening efforts have been reported over the last years [32–38]. However, the employed assay parameters cannot be directly applied to EHD proteins due to their weak ATPase activity. Therefore, developing a reproducible ATPase assay of EHDs compatible with HTS is crucial for potential drug screening efforts.

In this study, we have established the Malachite green (MLG)-based assay for measuring the liposome-stimulated ATPase activity of EHD4. MLG is a sensitive and cost-effective agent for determining released inorganic phosphate from ATP hydrolysis, and the assay does not include cumbersome washing or detection steps or the use of radioactive substances. Several assay parameters were optimized to allow HTS in the presence of liposomes. Eventually, we identified and initially characterized a prototypic small-molecule inhibitor for EHD4. The systematic optimization of this screen may be also applied to identify inhibitors for other dynamin-related proteins with a low, membrane-stimulated nucleotide hydrolysis activity.

## Materials and Methods

### EHD4 expression and purification

An N-terminal truncated construct of mouse EHD4 (Uniprot Q1MWP8, residues 22-541, EHD4^ΔN^) was expressed as N-terminal His_6_-fusion followed by a PreScission cleavage site in *E. coli* BL21 DE3 Rosetta (Novagen) from a modified pET28 vector, as described in [7]. Bacterial cultures were grown in Terrific Broth medium (TB) to an OD_600_ of 0.8 at 37 °C. Protein expression was induced at 18 °C by adding 100 μM isopropyl-ß-D-1-thiogalactopyranoside (IPTG) and cultures were grown for another 20 h at 18 °C. Cells were sedimented, resuspended in 50 mM HEPES pH 7.5, 500 mM NaCl, 25 mM imidazole, 2.5 mM β-mercaptoethanol, 2 mM MgCl_2_ and stored at –20 °C.

Upon thawing, 1 μM DNase (Roche), 500 μM of the protease inhibitor 4-(2-aminoethyl)-benzolsulfonyfluorid hydrochloride (BioChemica) and 5 U/mL^-^benzonase (Merck) were added (final concentrations). Cells were disrupted by passing them through a microfluidizer (Microfluidics). The lysed bacterial suspension was then centrifuged at 20,000 rpm for 30 min at 4 °C. The supernatant was collected and filtered using 0.45 µm filter packs, and then applied on a Ni^2+^-sepharose column equilibrated with equilibration buffer (50 mM HEPES pH 7.5, 500 mM NaCl, 25 mM imidazole, 2.5 mM β-mercaptoethanol, 2 mM MgCl_2_). This was followed by a long and short wash step with 50 mM HEPES pH 7.5, 700 mM NaCl, 30 mM imidazole, 2.5 mM β-mercaptoethanol, 1 mM ATP, 10 mM KCl and 2 mM MgCl_2_ and with equilibration buffer, respectively. Protein was eluted with 20 mM HEPES pH 7.5, 500 mM NaCl, 300 mM imidazole, 2.5 mM β-mercaptoethanol and 2 mM MgCl_2_, incubated with 1:150 (w/w) His-tagged Human Rhinovirus (HRV) 3C protease and dialyzed overnight in a 10 kDa cutoff membrane against dialysis buffer (20 mM HEPES pH 7.5, 500 mM NaCl, 2.5 mM β-mercaptoethanol and 2 mM MgCl_2_). To remove the protease and His_6_-tag, the cleaved protein was applied to a second Ni^2+^column equilibrated with equilibration buffer to which it bound also in the absence of the His-tag. Protein was eluted with 20 mM HEPES pH 7.5, 500 mM NaCl, 2.5 mM β-mercaptoethanol, 40 mM imidazole and 2 mM MgCl_2_ buffer. Cleaved protein was concentrated and loaded on Superdex 200 gelfiltration column (GE) equilibrated with size exclusion buffer (20 mM HEPES pH 7.5, 500 mM NaCl, 2.5 mM β-mercaptoethanol and 2 mM MgCl_2_). Fractions containing pure protein were pooled, concentrated and flash-frozen in liquid nitrogen and stored at –80 °C. Mouse EHD2 (Uniprot Q8BH64, residues 1-543) and human DNM1L isoform 2 (UniProtID: O00429-3, residues 1–710) were purified in a similar fashion [4, 39].

### Liposome preparation

Different lipid extracts were obtained from Sigma Aldrich with stock concentration of 25 mg/ml in chloroform. For small scale applications, 20 µl from the liposome stock was mixed with 200 µl of 3:1 (v/v) chloroform/methanol mixture in a glass vial. Lipids were dried under a gentle argon stream and stored overnight under vacuum in a desiccator. Dried lipids were hydrated for 5 min by adding 250 µl liposome buffer (20 mM HEPES pH 7.5, 300 mM NaCl) to reach a final concentration of 2 mg/ml and then resuspended by sonication in an ultrasonic water bath for 2 cycles of 30 s each. The freshly prepared liposomes were left for at least half an hour at room temperature before use. Prior to usage, the liposomes were extruded by passing 13 times through a mini-extruder (Avanti) using polycarbonate filters with 0.4 μm pore size. Larger volumes of liposomes were prepared as detailed in Fig. S1.

The following lipids were used in this study: Folch (Sigma-Aldrich, #B1502), brain-derived phosphatidyl serine (PS, Avanti-Polar Lipids, #840032C), synthetic 1-palmitoyl-2-oleoyl-sn-glycero-3-phospho-L-serine (POPS, Avanti-Polar Lipids, #840034P), synthetic 18:1 1,2-dioleoyl-sn-glycero-3-phospho-L-serine (DOPS, Avanti-Polar Lipids 840035).

### Malachite green dye preparation

0.1% MLG dye (Sigma, M9015) was dissolved in 1 M HCl by vortexing. 1% ammonium molybdate (Sigma, 09878-25G) was added and vigorously vortexed until no visible precipitates were observed. The resulting solution was filtered through a 0.45 μm filter and stored in an aluminum foil-wrapped glass bottle in the dark.

### ATPase assay

ATPase assays were performed in a 384 well plate at 25 °C (if not otherwise indicated). The assay buffer contained 20 mM HEPES, 150 mM KCl, 0.5 mM MgCl_2_, pH 7.5. 30 μM of ATP was mixed with 50 μg/ml DOPS in assay buffer. Reaction was started by adding 200 nM EHD4^ΔN^. At any given time-point, 25 μl of the reaction was taken out and mixed with 75 μl of the dye in a 384 well plate. The reaction gets quenched by the addition of the dye which denatures the protein. Absorbance at 650 nm was measured in a plate reader (Tecan, Infinite M200).

Turnover of substrate was limited to 10% for Michaelis-Menten constant determination (K_m_) and was calculated by fitting initial velocities to a non-linear least fit squares to the Michaelis Menten equation (v_0_ =v_max_[S]/(K_m_+[S])) using GraphPad Prism 7.05. ATPase assay and K_m_ determination were done in duplicates and triplicates, respectively.

### ATPase assay with HPLC

ATPase assay was performed as mentioned above and at defined time points, reaction aliquots were diluted three times in reaction buffer (50 mM Tris, 150 mM NaCl, pH 7.5) and quickly flash frozen in liquid nitrogen. Samples were applied to an HPLC and nucleotides were separated with reversed phase column (C18 100*4.6 mm) equilibrated with running buffer containing 100 mM potassium phosphate buffer pH 6.5, 10 mM tetrabutylammonium bromide, 7.5% (*v/v*) acetonitrile. Adenine nucleotides were detected by absorption at 254 nm and quantified by integrating the corresponding nucleotide peaks and determining the ADP/ATP ratios.

### Drug screening and IC_50_ determination

Screening was conducted at the Screening Unit of the Leibniz-Forschungsinstitut für Molekulare Pharmakologie (FMP) using a kinase drug library and the ‘diversity set’. 15,840 compounds were tested as singlets in 384 well plate format. A master mix (MM) was prepared in the assay buffer to have a final ATP concentration of 30 µM and DOPS concentration of 50 μg/ml. A 10 mM DMSO solution of the compounds was serially diluted, first with DMSO and then with the MM to a final concentration of 10 µM (1% of DMSO), using the Tecan-Evo workstation (pipetting robot). EHD4^ΔN^ (200 nM) was pipetted using multi-drop dispensing cassette. To have homogenous enzymatic activity, the plates were vortexed using a microplate mixer at 2,000 rpm for 10 s. The plates were incubated for 15 min at room temperature and 75 μl of MLG dye added with the cassette. The plates were measured in a Perkin Elmer Envision plate reader.

Data were analyzed with an automated pipeline [40] using KNIME software [41]. For the individual plates, the corresponding Ź values and SNR were calculated. Compared to the controĺs absorbance at 650 nm (Abs650), compounds were classified as inhibitors if the %Abs650 was ≤ 25%. These hits were picked from the drug library and used for further validation of IC_50_ determination and counter screening. IC_50_ values were determined in the same setup with varying inhibitor concentrations from 0.125 μM to 50 μM.

### Thermal shift assay (TSA)

TSA was done in a CFX96 touch-real time PCR using a fluorescent dye from ThermoFisher Scientific (catalogue number 4461146). Reactions (5 μM EHD4^ΔN^, 50 μM inhibitor and 1X dye) were prepared on ice, to a final volume of 50 μl and transferred to a 96 well PCR plate and then sealed with a plastic film. CFX96 was precooled to 4 °C and the PCR plate was inserted. The fluorescence was measured from 4 °C to 80 °C with 0.5 °C temperature steps. Protein melting temperature was calculated with the provided software (Bio-Rad) and plotted as bar diagram using GraphPad Prism 7.05.

### Compounds

The identified compounds are N-(2-hydroxyphenyl)-6-(3-oxo-2,3-dihydro-1,2-benzothiazol-2-yl)hexanamide (MS1, Molport, 008-333-699), N-(4-methyl-1,3-thiazol-2-yl)-3-phenoxybenzamide (MS8, Enamine, Z30820224) and 6-(Morpholino(4-(Trifluormethyl)phenyl)methyl)-1,3-Benzodioxol-5-ol (MS10, Enamine, Z99601030).

## Results

### Developing an EHD4 ATPase assay suitable for drug screening

We aimed for developing a prototypic small molecule inhibitor for the EHD family. From the four mammalian EHDs, only EHD2 and EHD4 can be bacterially expressed and purified in quantities compatible with HTS (Fig. S2A) [4, 7]. EHD2 has a very low basal ATPase activity (k_cat_ ∼1 h^-1^) which can only be moderately stimulated in the presence of lipids (k_cat_∼ 0.1 min^-1^) [4]. This very low activity is not suited for reliable HTS. Instead, we chose an EHD4 construct with a truncated N-terminus (EHD4^ΔN^), which was previously shown to have a higher basal and stimulated ATPase activity compared to EHD2 [7]. Since the basal EHD4^ΔN^ activity is still low, we focused on its liposome-stimulated ATPase activity.

To apply an MLG-based assay, the maximum absorption wavelength for the MLG dye in complex with orthophosphate (colorimetric complex) was initially determined. Various absorption maxima for MLG have been previously reported [42–45]. The dye used in this study showed an absorption maximum between 640 nm and 650 nm (Fig. S2B). Orthophosphate detection was linear up to 100 μM phosphate (Fig. S2C).

We next checked for interference of the MLG assay with assay components. ATP yielded a moderate background signal, maybe from phosphate contaminations, thereby restricting the maximal ATP concentration for screening (Fig. S2D, S2E). Furthermore, previously reported ATPase activities of EHD4^ΔN^ [4, 7] were recorded in the presence of Folch liposomes, which resulted in a large background signal in the MLG assay (Fig. 1A). In contrast, liposomes prepared from phosphatidyl serine (PS) from natural sources and synthetic 1,2-dioleoyl-sn-glycerol-3-phospho-L-serine (DOPS) gave a negligible background signal (Fig. 1A). However, only with synthetic DOPS, the EHD4^ΔN^ enzymatic activity was reproducibly stimulated (Fig. S2F, G). EHD4^ΔN^ enzymatic activity was higher at 30 °C and 25 °C compared to 37 °C (Fig. 1B, Fig. S2H). For easier application in the HTS assay, we used room temperature from hereon (25 °C).

**Fig. 1:**
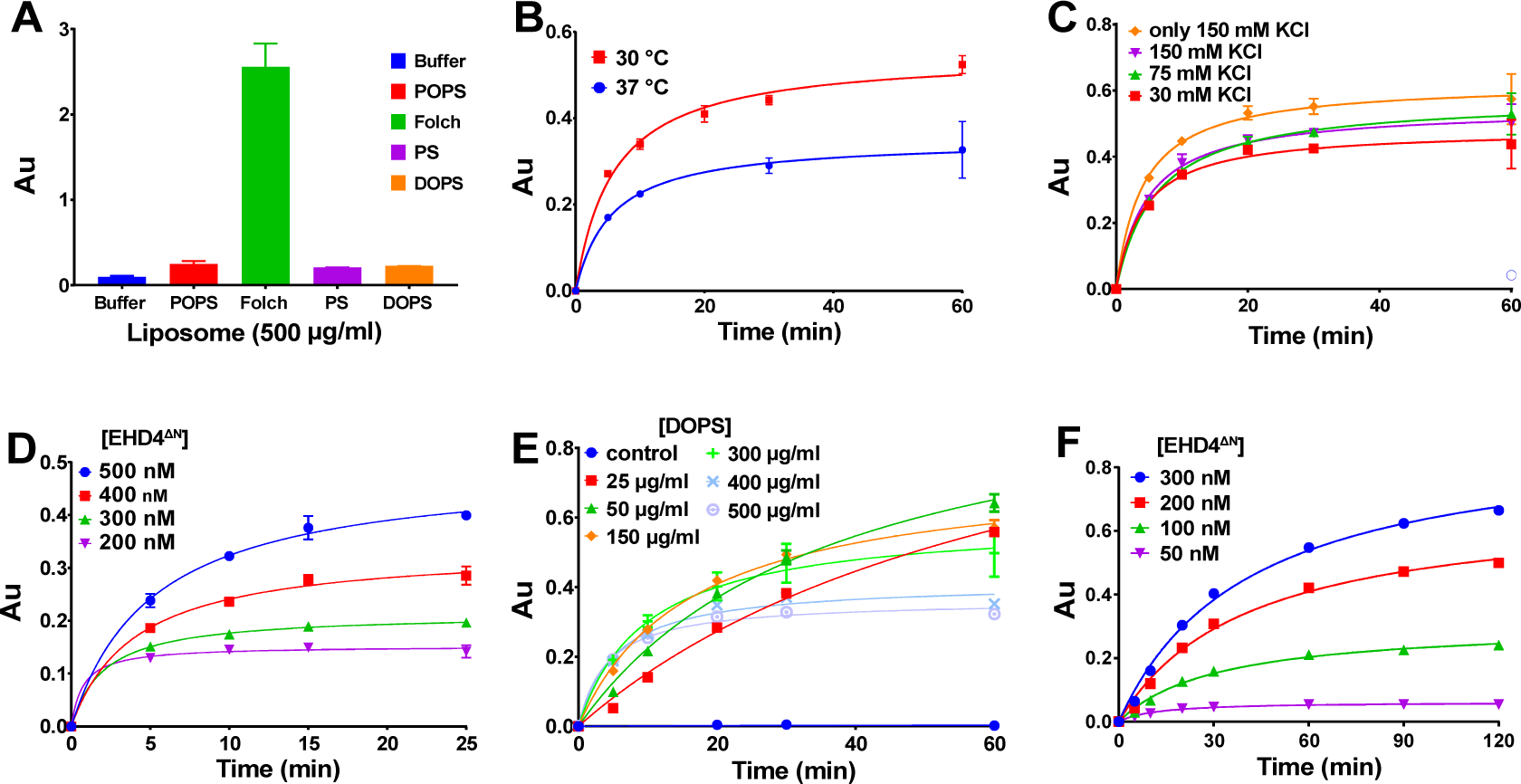
Optimization of the Malachite-Green ATPase assay for EHD4^ΔN^. **A** Background signal in the MLG assay arising from different lipids: Synthetic POPS, PS isolated from porcine brain and synthetic DOPS at 500 μg/ml. AU – arbitrary unit. **B** MLG-based ATPase assays for 4 μM EHD4^ΔN^, 500 μg/ml DOPS and 50 μM ATP were done at 37 °C and 30 °C. A further decrease to 25 °C did not affect the enzymatic activity compared to 30 °C (Fig. S2H). **C** MLG-based ATPase assays at 1 μM EHD4^ΔN^, 500 μg/ml synthetic DOPS and 30 μM ATP done at 25 °C in the presence of different concentrations of potassium ions. Removal of sodium ions in the presence of 150 mM KCl (denoted as ‘only 150 mM’) resulted in maximum ATPase activity. **D** EHD4^ΔN^ ATPase activity in the presence of 500 μg/ml DOPS and 30 μM ATP at 25 °C was screened at the indicated protein concentrations. **E** Dependency of the EHD4^ΔN^ ATPase activity on synthetic DOPS-liposome concentration was analyzed at 400 nM EHD4^ΔN^, 50 μM ATP in standard buffer at 25 °C. At lower DOPS-liposome concentration, an increase in enzymatic activity was observed. 50 μg/ml synthetic DOPS-liposomes were used in all subsequent assays. **F** Different EHD4^ΔN^ concentrations were tested in the presence of 50 μg/ml DOPS liposomes and 50 μM ATP at 25 °C. 200 nM EHD4^ΔN^ was chosen to have a SNR of at least 2. Data points represent the mean of two or more independent experiments and the error bar signifies the range of the measurements. When the range is smaller than the size of the data points, it is not displayed.

Potassium ions can influence the ATPase activity of various ATPases [46]. For dynamin and related enzymes, potassium has a catalytic role by binding into the active site next to the β-phosphate during GTP hydrolysis [47, 48]. We therefore tested the influence of potassium ions on EHD4^ΔN^ ATPases activity in addition to 150 mM NaCl already present in the assay buffer. In this case, 150 mM KCl resulted in the maximal activity (Fig. 1C). Replacing all sodium ions in the assay buffer with potassium led to a slight further increase in the ATPase activity (Fig. 1C). The optimized buffer conditions were therefore 20 mM HEPES (pH 7.5), 150 mM KCl and 0.5 mM MgCl_2_.

EHD4^ΔN^ ATPase assays were previously performed at μM protein concentrations which is too high for HTS carried out at typical inhibitor concentration of 10 μM. Therefore, we performed EHD4^ΔN^ titrations and found that EHD4^ΔN^ was active also at high nM concentrations at the improved assay conditions (Fig. 1D). Similarly, to analyze the liposome concentration dependency, we titrated different DOPS concentration at 400 nM EHD4^ΔN^ and found maximal activity at DOPS concentration of 50 μg/ml, e.g. 20-fold lower than previously described (Fig. 1E) [4, 7]. As protein and liposome concentrations mutually affect the enzymatic activity, we re-analyzed the ATPase dependence on EHD4^ΔN^ concentrations at a constant DOPS concentration of 50 μg/ml. A protein-dependent increase in ATPase activity was observed (Fig. 1F). To obtain a signal to noise ratio (SNR) of at least 2 while at the same time using a low protein concentration, 200 nM EHD4^ΔN^ and 50 μg/ml DOPS were chosen for the subsequent enzymatic assays.

### Enzyme kinetics and assay quality

For drug screening, we aimed at applying a substrate concentration close to the Michaelis-Menten constant (K_m_) to target competitive and non-competitive inhibitors [49]. The apparent K_m_ of EHD4^ΔN^ at a protein concentration of 1 µM was (20 ± 3) µM with a k_cat_ of 2.5 min^-1^ (Fig. 2A, B). However, due to the slightly higher SNR of 2.1 vs. 1.6 at 30 µM vs. 20 µM ATP after 15 min, respectively (Fig. 2C), we chose 30 μM ATP as substrate concentration for the subsequent HTS screens. The final assay parameters were 200 nM EHD4^ΔN^, 30 μM ATP, 50 μg/ml DOPS at 25 °C in 20 mM HEPES (pH 7.5), 150 mM KCl, 0.5 mM MgCl_2_ (Fig. 2D), which resulted in an excellent quality indicator for HTS (Ź-factor) [50] of 0.895.

**Fig. 2:**
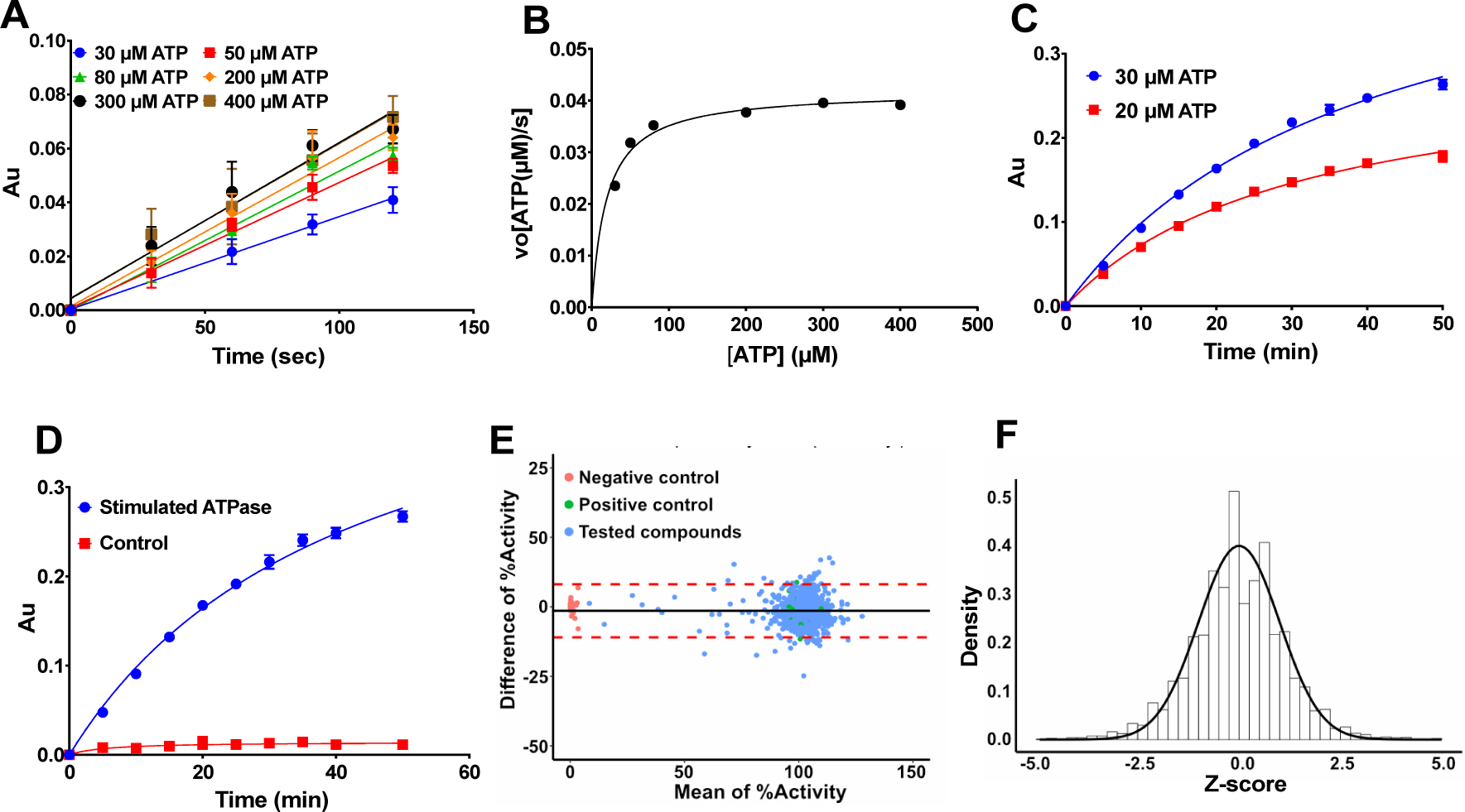
Reaction kinetics and assay quality. **A** Initial velocity of 1 µM EHD4^ΔN^ at different ATP concentrations in the presence of 50 µg/ml DOPS-liposomes were determined by calculating the slope of the linear reaction. **B** K_m_ was determined by first calculating the amount of hydrolyzed ATP in A using the standard curve mentioned in Fig. S2C and then plotting the initial rates of the reactions versus the substrate concentration. The kinetic parameters for EHD4^ΔN^ are K_m_ = (20 ± 3) μM, k_cat_ = (2.51 ± 0.07) 1/min, v_max_ = 2.51 μmoles ATP/min. **C** ATPase activity 200 nM of EHD4^ΔN^ in the presence of 50 µg/ml liposomes at 20 and 30 μM ATP concentration. The signal to noise ratio was 1.6 for 20 μM ATP whereas it was 2.1 with 30 μM ATP at 15 min. Therefore, 30 μM ATP was chosen for drug screening. **D** Ź in the final optimized assay condition (200 nM EHD4^ΔN^, 50 μg/ml DOPS and 30 μM ATP at 25 °C) was 0.895 at 15 min. Control represents assay without EHD4^ΔN^. **E** Bland-Altman plot from pilot screen using 3 library plates in technical replicates. Plot shows differences of repeated measurements falling into a 95% confidence interval of ± 9.5%, suggesting high confidence between replicates. **F** From the same pilot screening, a Z-score histogram was plotted showing normal distribution. Data points represent the mean of two or more independent experiments and the error bar signifies the range or standard deviation. When the range/standard deviation is smaller than the size of the data point, it is not displayed.

To visually assess data reproducibility of our assay, we performed a pilot screen using three plates from the drug library, measured in two technical replicates. Data were analyzed using the Bland-Altman method (Fig. 2E) by plotting the difference of percentage activities of the two technical replicates against their mean. The coefficient of repeatability was calculated from the observed differences and showed a high confidence between the replicates, with the differences of repeated measurements falling into a 95% confidence interval of ±9.5%. In a Z-score histogram (Fig. 2F), Z-scores were equally distributed around the zero midpoint, with a shape resembling a normal distribution, indicating that data normalization was effective in removing the batch-to-batch variation in absolute signals. Ź-factors of the plates were between 0.75 – 0.9. Finally, no effect of 1.5% DMSO on the enzymatic activity was observed (Fig. S2I). This rendered the assay compatible for the HTS setup, since all small molecules in the HTS were diluted from DMSO stock solutions, yielding a final DMSO concentration in the assay of 1%.

### Drug screening

Using the optimized assay conditions, we screened ∼16,000 compounds from the drug library of Leibniz-Forschungsinstitut für Molekulare Pharmakologie (FMP) for EHD4^ΔN^ ATPase inhibition in single replicates. Data were analyzed with an automated pipeline [40]. Compounds showing a decrease in enzymatic activity by 25% were grouped as inhibitors. Primary hits were validated in duplicates and counter-screened for false positives. For counter screening, we tested 30 μM phosphate in assay buffer against 10 μM inhibitor in the MLG assay to determine if inhibitors interfered with the colorimetric reaction. IC_50_ values for the validated hits were determined. An overview of the drug screening and a scatter plot of the single point screening data is shown in Fig. 3A and B.

**Fig. 3:**
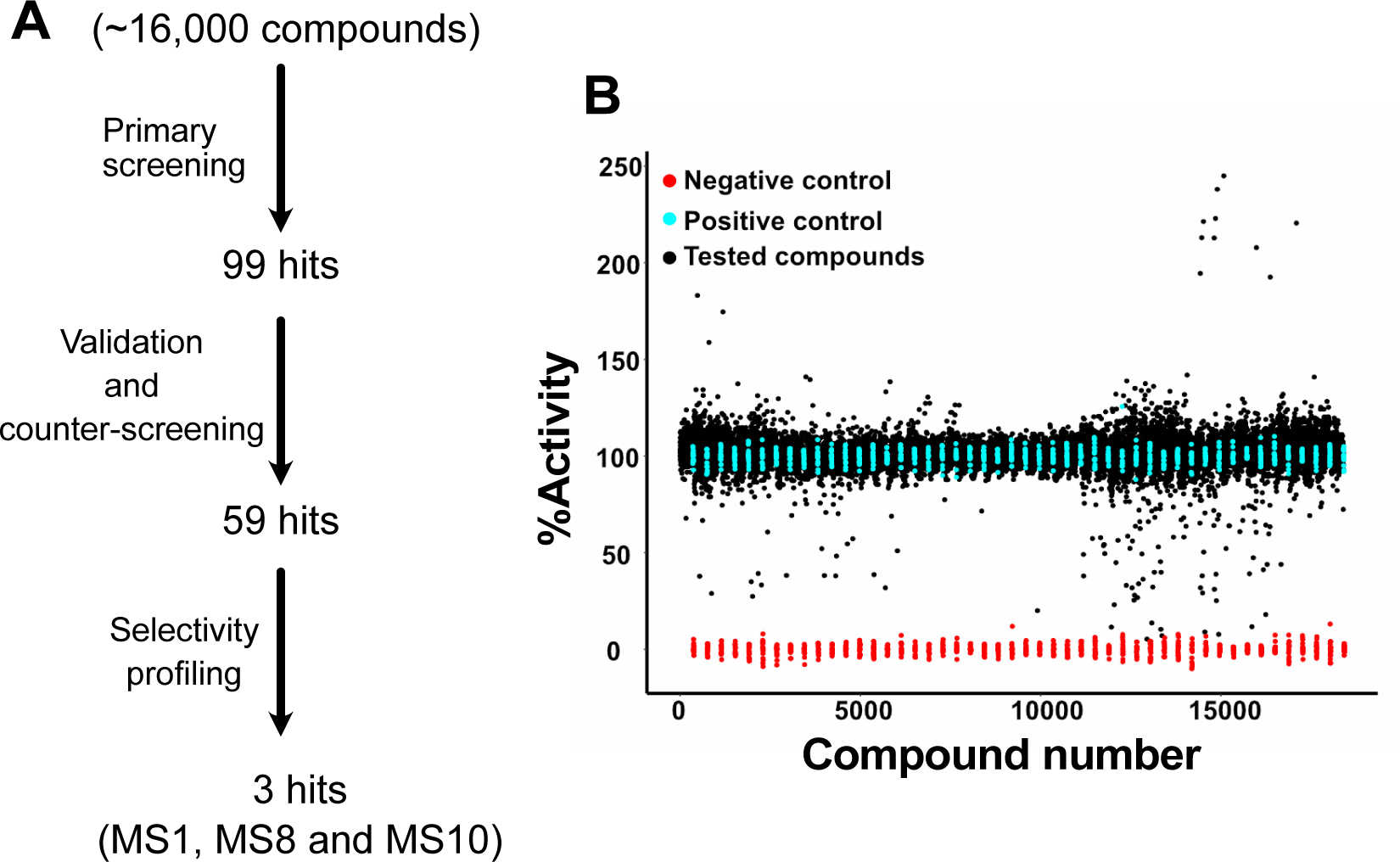
EHD4 drug screening. **A** Schematic representation of EHD4 drug screening. Approximately 16,000 compounds were tested at 10 μM concentration in a primary screen against EHD4^ΔN^ with the MLG assay. Initial hits were validated in duplicates and counter-screened to eliminate false positives, e.g. compounds that interfered with the colorimetric reaction. The final three compounds as primary hits from drug screening were selected based on their IC_50_ value and chemical structure. Thus, compound with an unstable chemical structure prone to oxidation or degradation, with a tendency to form covalent bonds, and containing a PAINS substructure were eliminated, as well as compounds showing a steep fall in IC_50_ curves (see Discussion). **B** Scatter plot of the single point screening data. Compound numbers are for visualization purposes. Cyan represents the positive controls without compounds, red the negative controls without EHD4^ΔN^, and black the tested compounds. Potential activators, e.g. compounds apparently showing a higher activity, were all later found to be false positives in a counter screening experiment by inducing themselves a signal in the MLG assay.

For the final compound selection, we considered IC_50_ values and excluded chemical structure of compounds that were unstable and hence prone to degradation. Further selection criteria are outlined in the discussion. We proceeded with three primary hits, MS1, MS8 and MS10, showing IC_50_ values of 0.92 μM, 3.8 μM and 2.9 μM respectively (Fig. 4), with maximal inhibition of 60-80%.

**Fig. 4:**
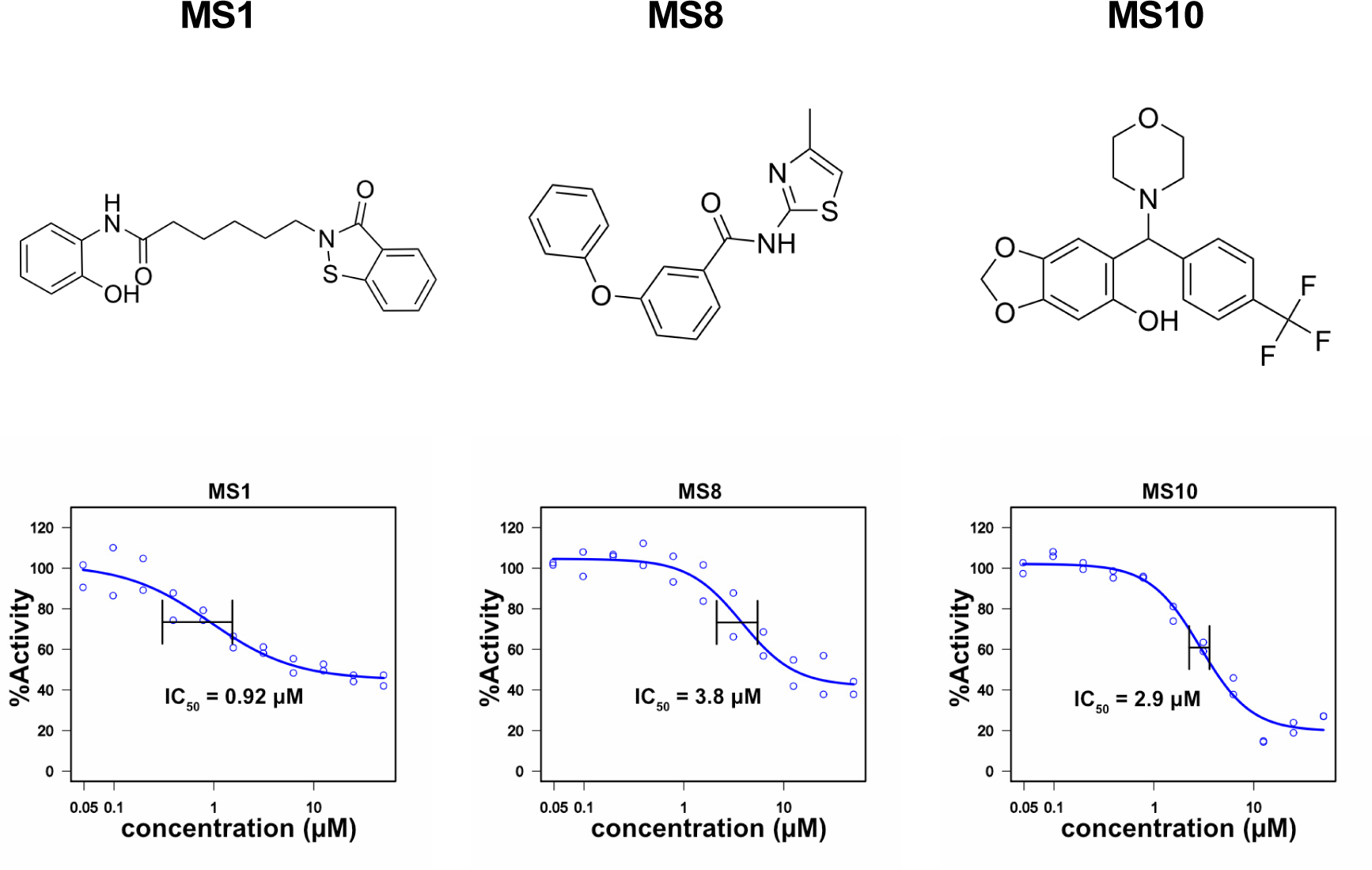
Primary hits from drug screening. Chemical structure and IC_50_ values for the primary hits from the EHD4 drug screen. Hits were picked from the drug library and IC_50_ values were determined in duplicates against an inhibitor concentration range from 0.125 μM to 50 μM.

### Biochemical characterization of the inhibitors

We re-purchased the three compounds and validated them first via the MLG assay (Fig. 5A) and then via HPLC to remove any assay bias (Fig. 5B). All three compounds inhibited EHD4^ΔN^ enzymatic activity in both assays. IC_50_ measurement for the purchased compounds (Fig. S3) were similar to the compounds picked from the drug library (Fig. 4). To examine specificity, we investigated the inhibitors against EHD2. Due to the exceptionally low ATPase activity of EHD2 and the high background signal by Folch lipids required for its ATPase stimulation, detection in the MLG assay was not possible. Thus, EHD2 ATPase assay was instead measured by HPLC (Fig. S4D) [4]. MS1 and MS10 inhibited EHD2 enzymatic activity whereas MS8 had no effect (Fig. 5C). In thermal shift assay, EHD4^ΔN^ showed a melting temperature of 44.5 °C (Fig. S2J). MS1 had a destabilizing effect on EHD4^ΔN^, as evident by a negative ΔT_m_ of –5 °C. MS10 also diminished EHD4^ΔN^ stability whereas MS8 had no such effect (Fig. 5D, Fig. S2K). MS8 was further counter-screened against a distantly related dynamin family member, DNM1L, by the MLG assay (Fig. S4A-C). MS8 showed no effect on GTPase activity of DNM1L (Fig. 5E) further validating its specific inhibition of EHD4. These control experiments also excluded a direct effect of MS8 on liposome integrity, required for the functional activation of EHD2 and DNM1L.

**Fig. 5:**
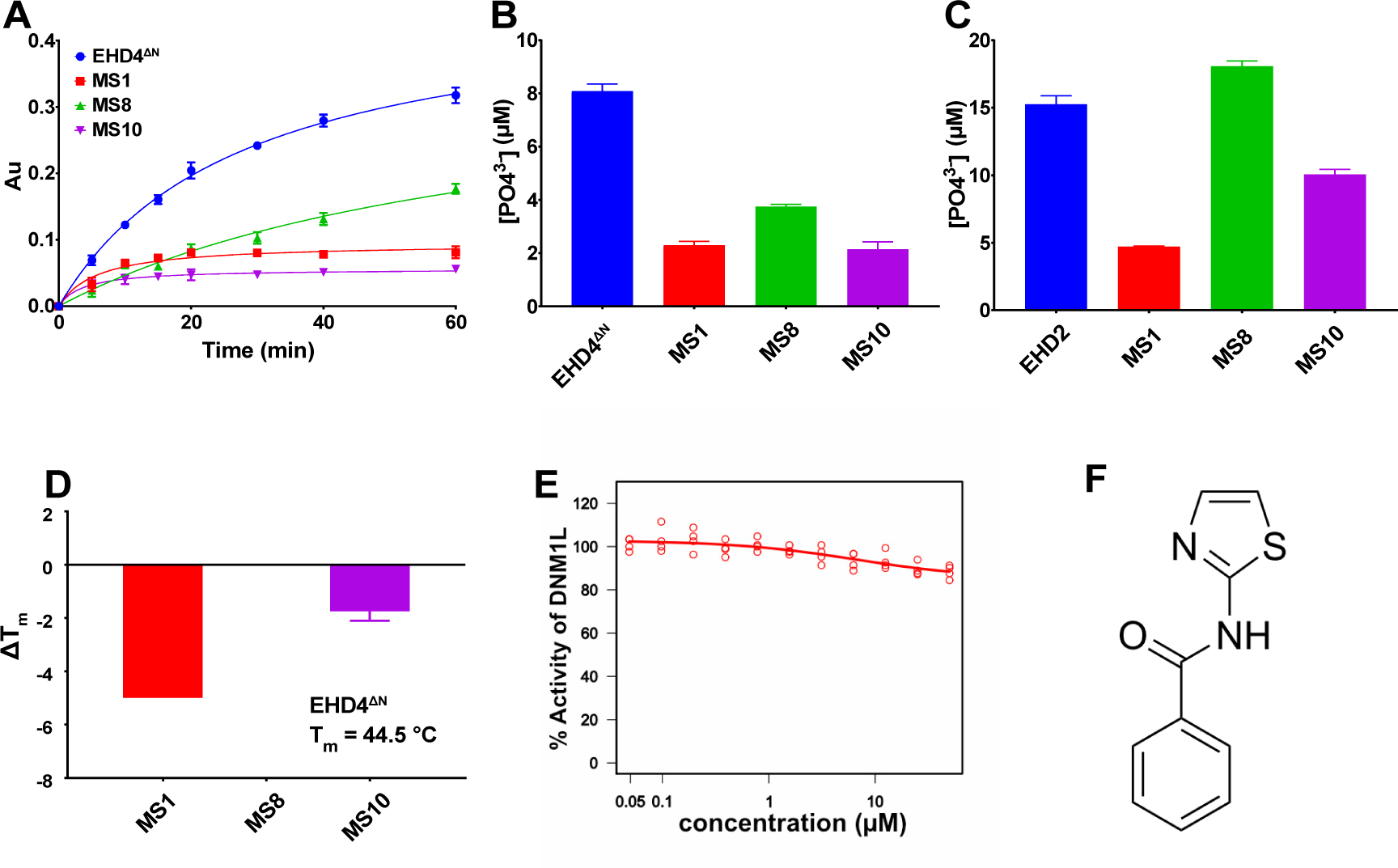
Biochemical characterization of the primary hits. **A** One-hour time course of the MLG-based enzymatic ATPase assay in the presence of 50 μM inhibitor. EHD4^ΔN^ (blue curve) represents the positive control. **B** Primary hits were validated at 50 μM inhibitor concentration in an HPLC setup to exclude assay bias. ATPase assays done in A and B were in standard conditions. **C** Primary hits were tested against EHD2. Only MS8 had no effect on EHD2 ATPase activity. ATPase assay was done at an EHD2 concentration of 5 μM in the presence of 500 μg/ml Folch liposomes and 50 μM ATP in assay buffer at 30 °C. **D** Changes in melting temperature of 5 μM EHD4^ΔN^ were studied in the presence of 50 μM inhibitor by thermal shift assay. MS1 and MS10 destabilized the protein, as evident by negative ΔT_m_, whereas MS8 had no effect on the stability of EHD4^ΔN^. **E** MS8 did not inhibit the GTPase activity of DNM1L. Shown is an IC_50_ curve plotted with an inhibitor concentration range from 0.125 μM to 50 μM. **F** MS8 substructure obtained by dissecting the diphenlyether bond. This substructure was used in our chemical search to have the highest diversity of the chemical space and the optimal coverage of chemical vectors to gain a structure-activity relationship. Data points represents the mean of two or more independent experiments and the error bar signifies the range or standard deviation. When the standard deviation is smaller than the size of the data point, it is not displayed.

### Structure activity relationship

Based on the chemical structure of the hit MS8, a substructure search of the FMP library was conducted. Two possible dissections of chemical bonds, either the amide or diphenylether bond, have been used in this search. The resulting fragments were analyzed according to the number of compounds inside the FMP library with highest diversity of the chemical space and the optimal coverage of chemical vectors to gain a structure activity relationship. In this process, the N-(thiazol-2-yl)benzamides substructure (Fig. 5F) yielded the best results of a suitable compound set containing this fragment.

Preliminary SAR data were derived by commercially available 55 compounds from the substructure search of N-(thiazol-2-yl)benzamides in the FMP library with a broad spectrum of substitution pattern. Only three compounds (Z5, Z7 and Z8, see Fig. S5) showed 20% or more inhibition of EHD4^ΔN^ ATPase activity in the MLG assay at 50 μM inhibitor concentration and were classified as active, whereas the remaining were classified as inactive (see Table S1).

Analysis of the SAR substructures showed that a connection between the benzamides to the thiazole rings in 2-position is mandatory for the activity. The substitution in 3-position of the thiazole ring can be either alkyl chains or aromatic sided chains (see Fig. S5 and Table S1, MS8, Z5 and Z7). Substitution in the 5-position diminishes the activity significantly (see Table S1, e.g. compounds 106268 or 106333). Any annealed rings on the benzamide or between the 4 and 5 position of the thiazole ring abandoned the activity completely (see Table S1, e.g. compounds 105229, 201598 or 203985 and many more). The benzamide ring does not tolerate substitutions in *ortho*-position (Table S1, compounds 209726 or 301620). Activity is gained via substitution in *meta*-position with methoxy or phenoxy rings only (Fig. S5 and Table S1, MS8, Z5, Z7 and Z8). Any other substitution with smaller functional groups in *para* and/or *meta*-position seems to diminish the activity (Table S1, e.g. compounds 110741, 100723 or 209726), but we have only a limited number of examples. In the future, efforts will be made to re-purchase commercial available N-(thiazol-2-yl)benzamides based on the knowledge gained in the summarized SAR (Fig. 6) in order to further boost the inhibitory activity of EHD4.

**Fig. 6:**
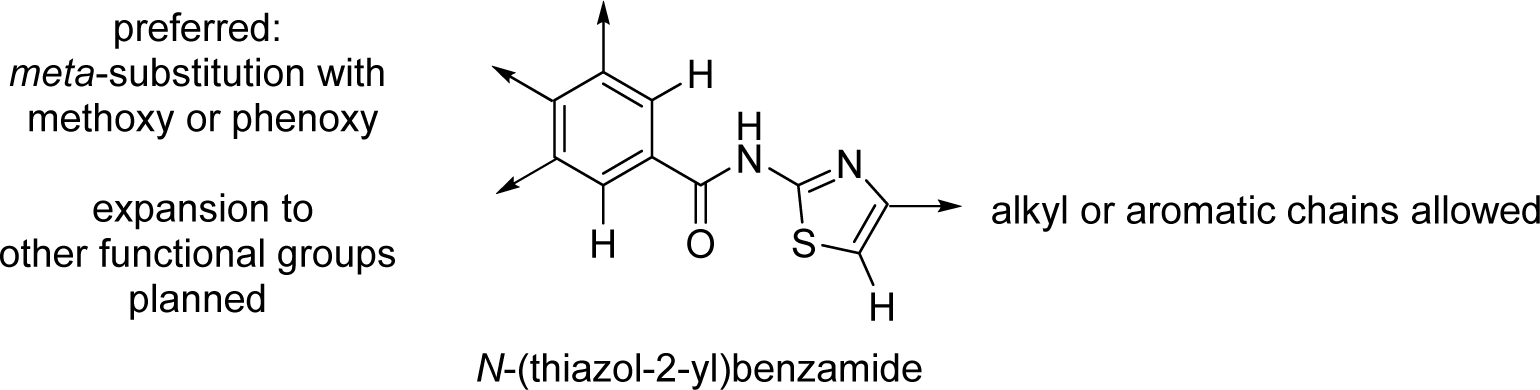
Initial SAR data on commercially available compounds tested in the EHD4 assay based on N-(thiazol-2-yl)benzamides as the central core scaffold. Substructure search resulted in 55 commercially available compounds which were picked from the drug library and re-tested. Only three of them inhibited EHD4 enzymatic activity by 20% or more (Fig. S5).

## Discussion

There are no chemical inhibitors for EHD proteins described to date. This study reports the identification of an EHD4 inhibitor by establishing a robust, reproducible EHD4 ATPase assay in the presence of liposomes, which is compatible with drug screening. For optimization of the MLG-based assay, the use of synthetic lipid, DOPS, played a major role. The chemical composition of natural extracts (Folch, PS or similar lipids) inevitably varies from batch to batch, and the extracts may include trace amounts of other lipids, which lead to variability in the enzymatic protein activity. Synthetic DOPS (18:1) used in this study has similar properties to naturally occurring brain PS, but it is more stable towards oxidation, which makes it a suitable substitute. DOPS has been used in drug discovery studies for other purposes, for example as a component of lipid mixtures to mimic the plasma membrane [51], in liposome preparations for anti-tumor applications [52] and amyloidosis studies [53].

Utilizing the MLG-based assay, we screened ≈16,000 compounds for their ability to inhibit EHD4^ΔN^ ATPase activity. After validating and counter screening, we excluded unstable chemical structure of compounds prone to degradation and or oxidation and potential covalent bonds forming compounds (Michael acceptor) that could lead to non-specific cellular binding [54]. We also removed substances inducing a sudden concentration-dependent activity drop of EHD4^ΔN^, which may hint at the compound forming aggregates and sequestering the enzyme [55, 56]. Furthermore, we applied substructure filtering for identifying Pan Assay Interference Compounds (PAINS) [57] and removed compounds showing high number of reported biological activities in the ChEMBL database [58]. In this way, we limited the further analysis to three compounds MS1, MS8 and MS10, which inhibited EHD4^ΔN^ with IC_50_ values in the low micromolar range. The preference for MS8 was guided by structural considerations and specificity towards the target. Although not automatically excluded via PAINS filter, MS1 consists of an isothiazolinone substructure which was listed as a nuisance compound in a GSK panel list [59]. Additionally, it is described that the structure might degrade over time via oxidation to sulfoxide and sulfone following by ring-opening [60]. MS10 might be synthesized via a mannich-like condensation mechanism using the corresponding ketone and the morpholine to the imine followed by reductive amination. One could consider a retro-condensation mechanism which would lead to degradation of the MS10. In comparison to MS1 and MS10, the structure of MS8 is very stable consisting of two aromatic groups connected via an amide bond. Moreover, MS8 did not inhibit enzymatic activity of the closely related EHD2 and also had no effect on the enzymatic activity of another dynamin superfamily protein member, DNM1L.

The exact inhibition mechanism of the compounds requires further analyses. Substances directly interfering with ATP binding, membrane binding or oligomerization would be expected to reduce stimulated ATP hydrolysis reaction. Also, allosteric inhibition of the ATPase could be envisaged. Furthermore, compounds stabilizing the EHD oligomers may lead to interference of catalysis, e.g. by blocking ADP release after one round of hydrolysis. A structure of an EHD4-inhibitor complex may provide details of the inhibition mode, but so far, we were not able to obtain a crystal structure of such a complex so far.

MS8 could be used to better understand the EHD4-mediated signaling pathways. EHD4 was discovered as a protein induced by nerve growth factor (NGF) and mediates cytoplasmic signaling of NGF through its receptor, TrkA (Tropomyosin receptor kinase A) [21]. Other roles of EHD4 include internalization of the trans-membrane protein Nogo-A into neuronal cells by EHD4-mediated endocytosis. Nogo-A is expressed in the adult central nervous system (CNS), where it inhibits axonal growth, regeneration and plasticity [61]. EHD4 along with EHD1 is also involved in the endocytosis of L1/neuron-glia cell adhesion molecule (NgCAM) in neurons known to regulate axonal growth [62]. Although progress has been made in addressing the specific role of EHD4 in this membrane trafficking pathway, several open questions remain. For example, it is unclear whether single or multiple routes for EHD4-mediated membrane trafficking exist [63]. Furthermore, whether other EHD proteins are also involved in co-regulating endocytosis for TrkA still needs to be addressed. MS8 may help to dissect EHD4’s cellular function. In particular, it will be interesting to explore whether MS8 shows an effect on endosomal signaling in neurons. If such an effect would be observed, even possible therapeutic applications could be considered. For example, a role for a more potent MS8 derivative could be to suppress Nogo-A signaling, which may help in functional recovery of the adult CNS in the aftermath of an injury by stimulating regeneration and nerve fiber growth. Such function would be supported by previous studies showing that EHD4 ATPase-deficient mutant completely block the internalization of Nogo-A [61].

A combined approach of HTS and structure-guided drug design could result in a highly selective potent EHD4 inhibitor. Our initial SAR studies have indicated preferred chemical modification of MS8 in order to develop more potent EHD4 inhibitors in the future.

## Authors Contribution

M.S. planned, performed and analyzed all experiments if not otherwise indicated. A.O. assisted with performing drug screening and IC_50_ measurement. M.N. performed the drug screening analyses and calculated IC_50_ with automated pipeline at FMP. E.S. carried out the hit picking and substructure search and SAR analysis. O.D. supervised the study. M.S. and O.D. wrote the manuscript with input from all the authors.

## Acknowledgements

We thank Dr. Marc Nazaré, Dr. Peter Lindermann and Jerome Paul for usage of and support with their Rotavac and Regina Piske and Helmut Kettenmann for granting continuous access to their plate reader. We thank the online scientific community for intense discussions, especially Dr. Adam Shapiro (Entasis Therapeutics), and Marius Weismehl for critical reading of the manuscript. We thank Deutsche Forschungsgemeinschaft (SFB958, project A12, to O.D.) and MDC for their funding and support.

## Supplemental Data

**Fig. S1:**
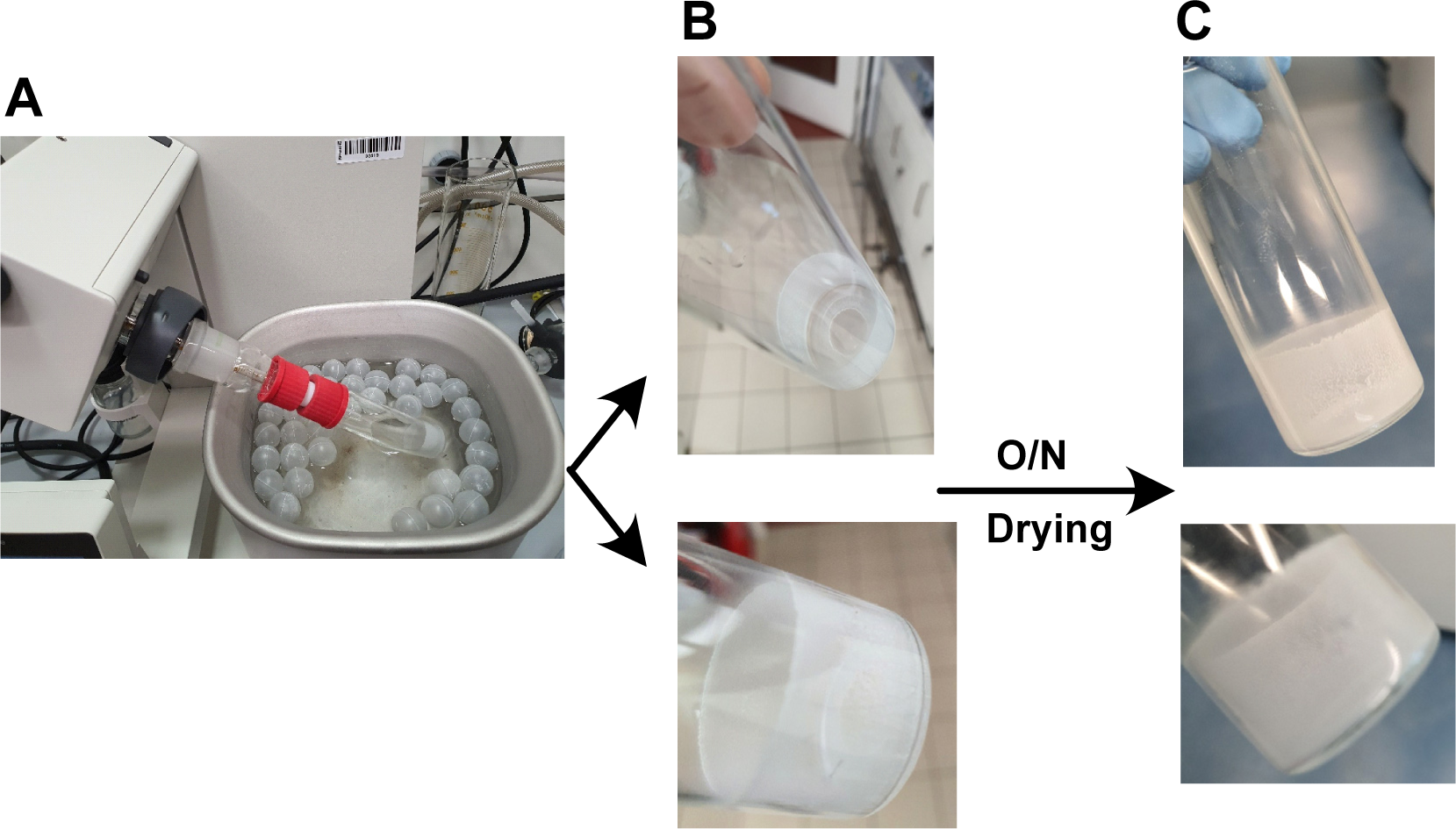
Schematic representation of liposome preparation with the help of a rotary evaporator. **A** Overview of the experimental setup. Nitrogen gas instead of vacuum was used to evaporate the chloroform/methanol mix from the lipid solution. For this, the vacuum port was removed and a 50 ml serological pipette attached to a nitrogen gas tubing into the glass vial (VWR 548-0156). 500 μl of DOPS (Avanti Polar Lipids) were mixed with 6 ml of a chloroform/methanol mixture (3:1 v/v) in the previously mentioned glass vial. This vial was attached to a Rotavapor R-300 (Buchi) with the bottom part of the vial dipping into the heating bath kept at 25 °C. A gentle nitrogen stream was introduced into the vial and adapted to not observe ripples on the surface of the solution. The rotation speed was set to 175 rpm. **B** Approximately 25 min later, a regular lipid monolayer formed on the glass vial after removal of chloroform/methanol mixture using the Rota vapor. **C** Same lipids after overnight (O/N) drying.

**Fig. S2:**
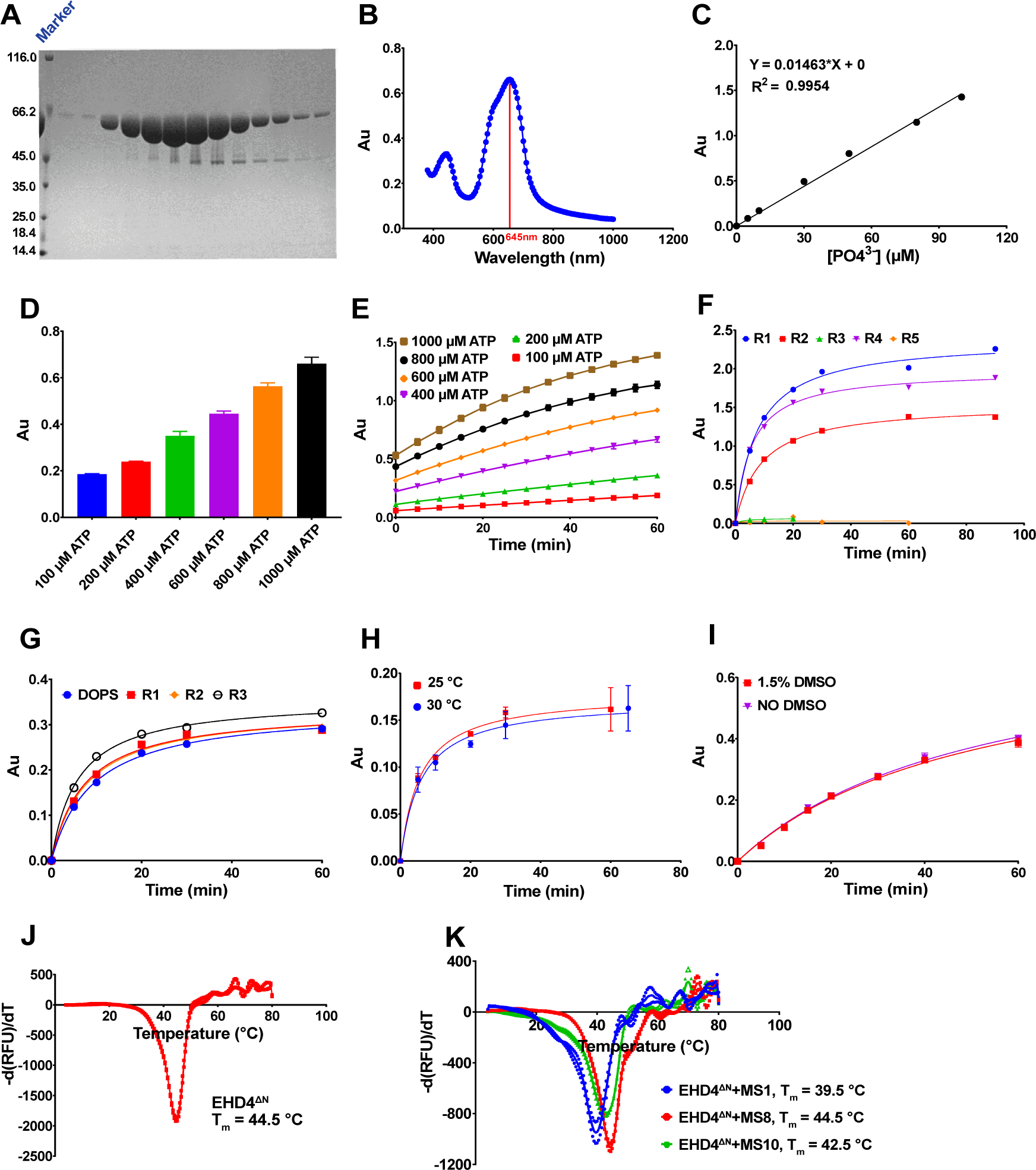
EHD4^ΔN^ ATPase assay optimization. **A** Coomassie-stained SDS-PAGE gel showing fractions of the final gelfiltration peak of the EHD4^ΔN^ purification. **B** Absorption maxima of the colorimetric complex formed between the MLG dye, molybdate and orthophosphate. AU – arbitrary units. **C** Standard curve of orthophosphate in the MGL assay. The determined fit parameters of the curve are shown on the top left. **D** Background signal from ATP possibly derived from phosphate contaminations is directly proportional to the ATP concentration in the assay. **E** ATP hydrolysis in the absence of EHD4 ^ΔN^ and DOPS in the acidic environment of the MLG dye over time. Note that higher ATP concentration leads to a higher background signal. **F** ATPase activity of EHD4^ΔN^ at 25 °C at 2 μM EHD4^ΔN^, 200 μM ATP and 300 μg/ml liposomes composed of natural PS in assay buffer was detected by the MLG assay, but it was not reproducible. R1-R5 represents different repetitions of the EHD4 assay under supposedly identical conditions. **G** ATPase assay of EHD4^ΔN^ at 4 μM EHD4^ΔN^, 30 μM ATP and 500 μg/ml synthetic DOPS liposomes in assay buffer were conducted at 25 °C. R1, R2 and R3 represent three independent preparations of DOPS liposomes from different batches, resulting in similar activities. **H** ATPase assay comparison at 30 °C and at 25 °C at a concentration of 20 μM ATP and 4 μM EHD4^ΔN^ and 500 μg/ml DOPS liposomes in assay buffer. **I** Inclusions of 1.5% DMSO had no effect on EHD4^ΔN^ enzymatic activity in the presence of liposomes, rendering the assay compatible with the HTS setup. **J** Thermal shift assay of EHD4^ΔN^ at a protein concentration of 5 μM to determine the melting temperature of the protein. **K** Thermal shift assay of 5 μM EHD4^ΔN^ with 10 μM each of MS1, M8 and MS10 to determine the melting temperature of the protein in presence of the inhibitors. Except for B and F, data points represent the mean of two or more independent experiments and the error bar signifies the range or standard deviation. When the standard deviation is smaller than the size of the data point, it is not displayed.

**Fig. S3:**
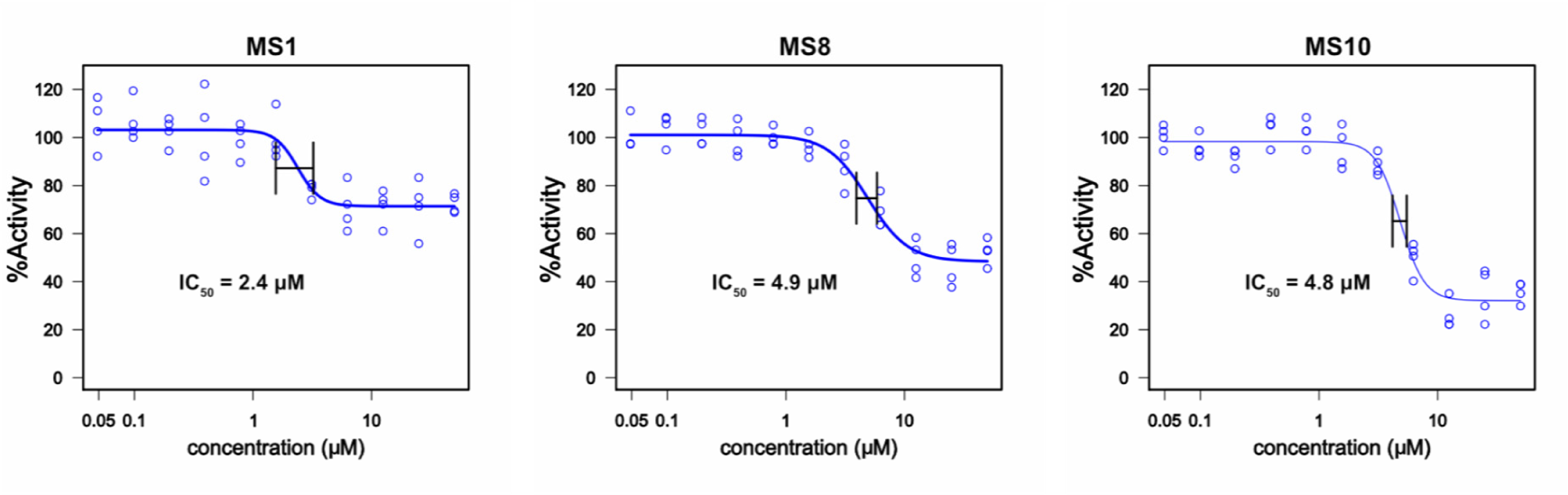
IC_50_ for the purchased primary hits. IC_50_ values for the purchased primary hits were similar to the previously determined IC_50_ values of the compounds from the drug library. Repurchased MS1 showed an IC_50_ value of 2.4 μM (vs 0.92 μM with the compound from the drug library), MS8 an IC_50_ value of 4.9 μM (vs 3.8 μM with the compound from the drug library), MS10 an IC_50_ value of 4.8 μM (vs 2.9 μM with the compound from the drug library).

**Fig. S4:**
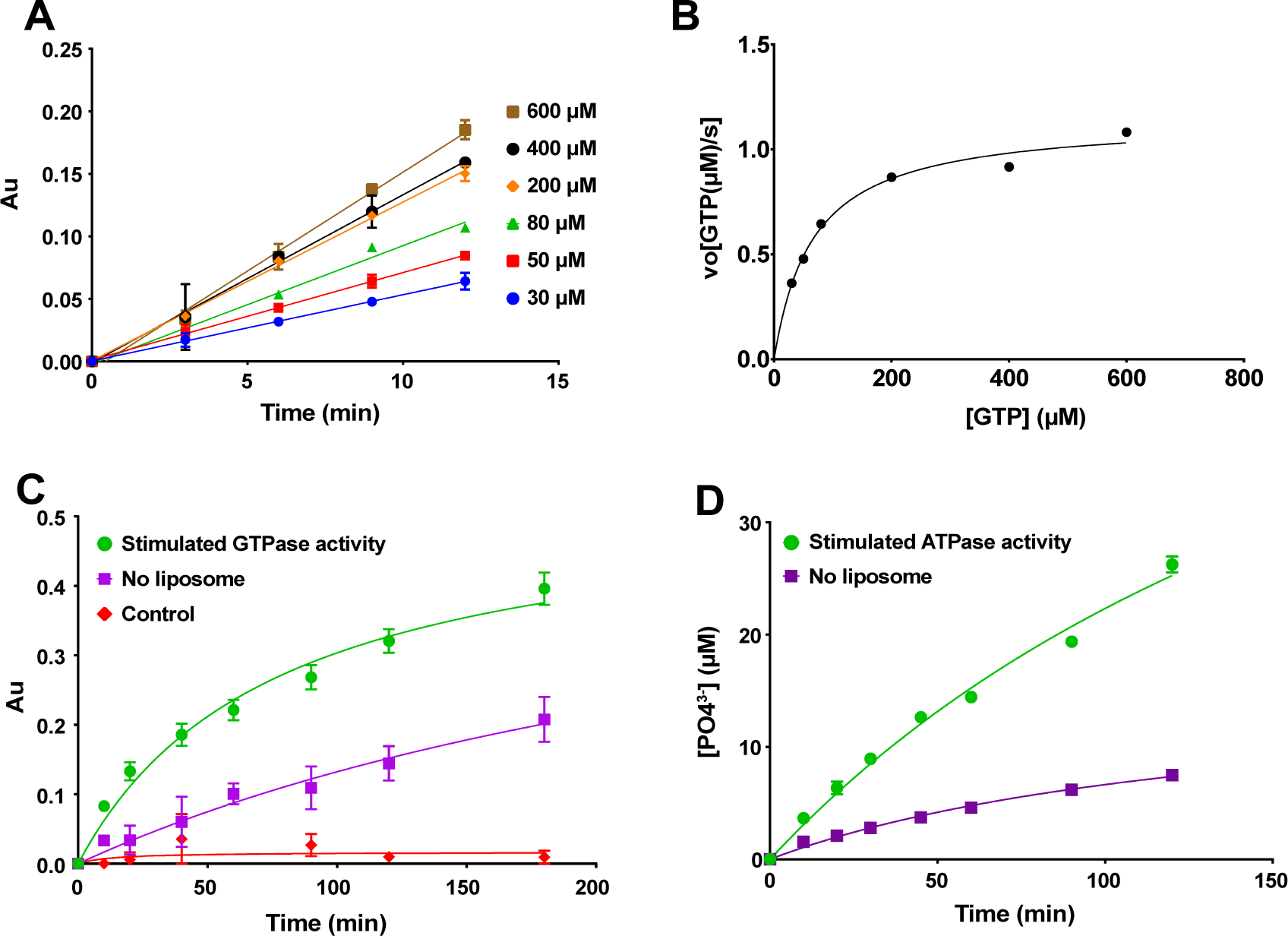
DNM1L and EHD2 enzymatic assay. **A** Initial velocities of DNM1L GTPase activity at 300 nM DNM1L, and, 500 μg/ml synthetic DOPS, T=25 °C and at different GTP concentrations were determined by a linear fit. **B** K_m_ was determined by first calculating the amount of hydrolyzed GTP in **(A)** using the standard curve mentioned in Fig. S2C and then plotting the initial rates of the reactions versus the substrate concentration. The kinetic parameters for DNM1L are K_m_ = (66 ± 9) μM, k_cat_ = (3.81 ± 0.15) 1/s, v_max_ = 1.15 μmoles GTP/s. **C** GTPase assay monitoring DNM1L activity were done with the final ptimized parameters, which were 300 nM DNM1L, 40 μM GTP, 200 μg/ml synthetic DOPS liposomes, T = 25 °C in 20 mM HEPES (pH 7.5), 150 mM KCl, 0.5 mM MgCl_2_. Ź and SNR were 0.72 and 2.1 at 20 min for the stimulated GTPase activity. **D** EHD2 ATPase activity by an HPLC-based method. Assay conditions were 5 μM EHD2, 50 μM ATP and 1 mg/ml Folch liposomes at 30 °C. Folch liposomes were used here as the HPLC-based assay is compatible with Folch liposomes. The data points except in B represent the mean of two independent experiments and the error bar signifies the range of the fit. When the standard deviation is smaller than the size of the data point, the error bar is not displayed.

**Fig. S5:**
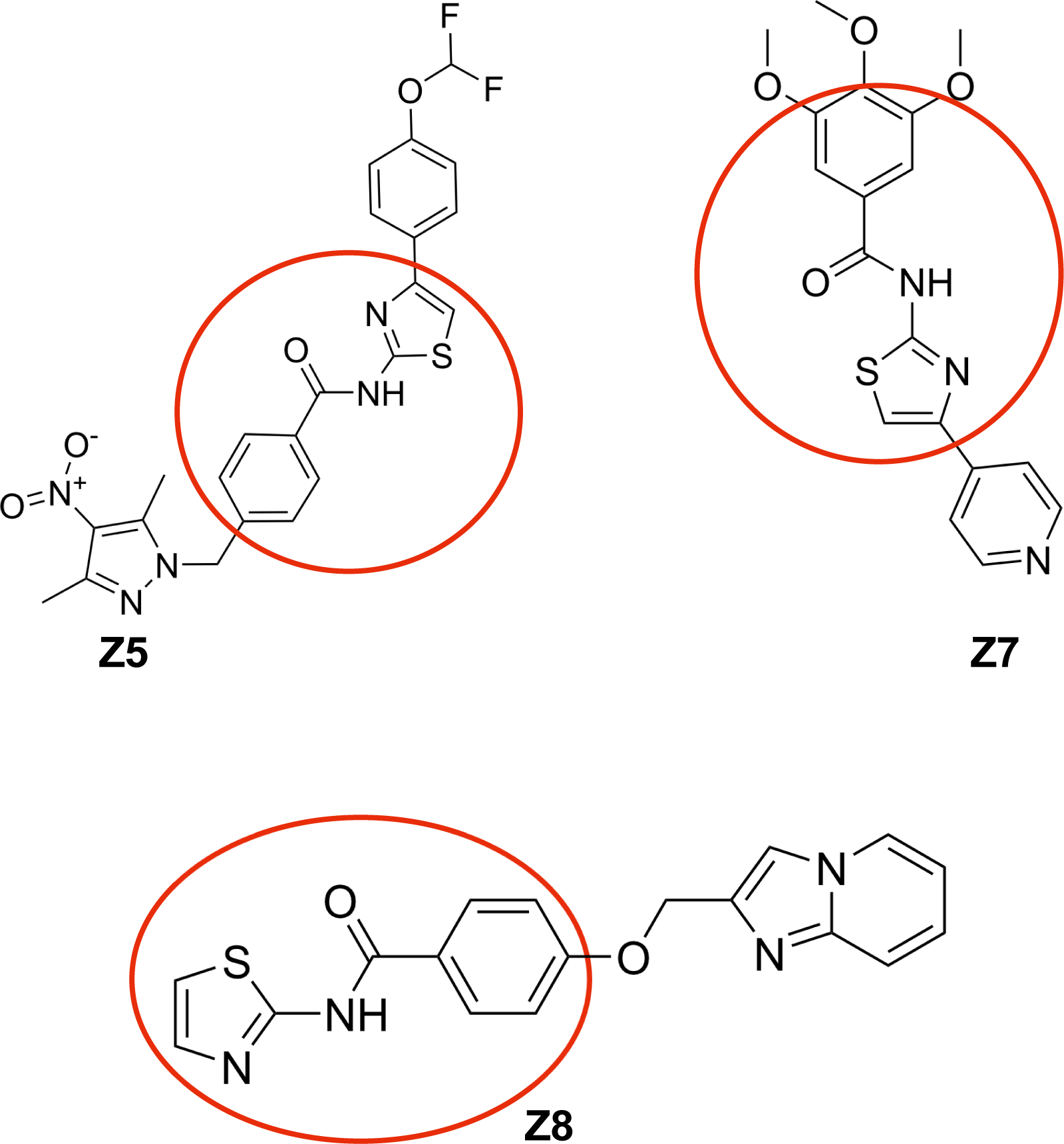
Compound structure of the Z5, Z7 and Z8. The three compounds that showed inhibition towards EHD4^ΔN^ at 50 μM inhibitor concentration by 20% or more in the MLG assay, among the 55 compounds from the substructure search in our SAR study. Z5, Z7 and Z8 inhibited EHD4^ΔN^ enzymatic activity by 21%, 44% and 38% respectively. The N-(thiazol-2-yl)benzamide moiety, used for our chemical search is encircled in red.

**Table S1:**
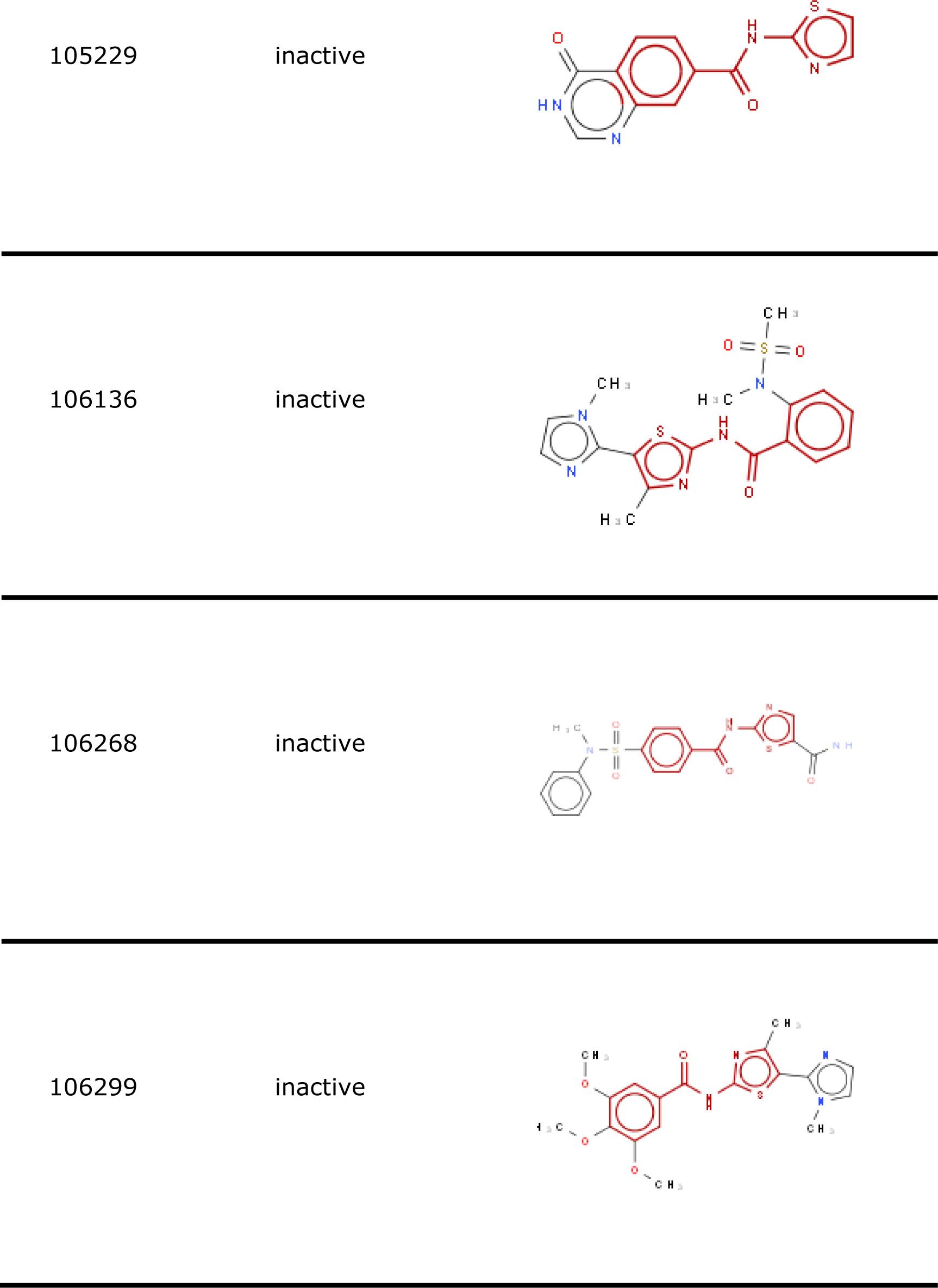
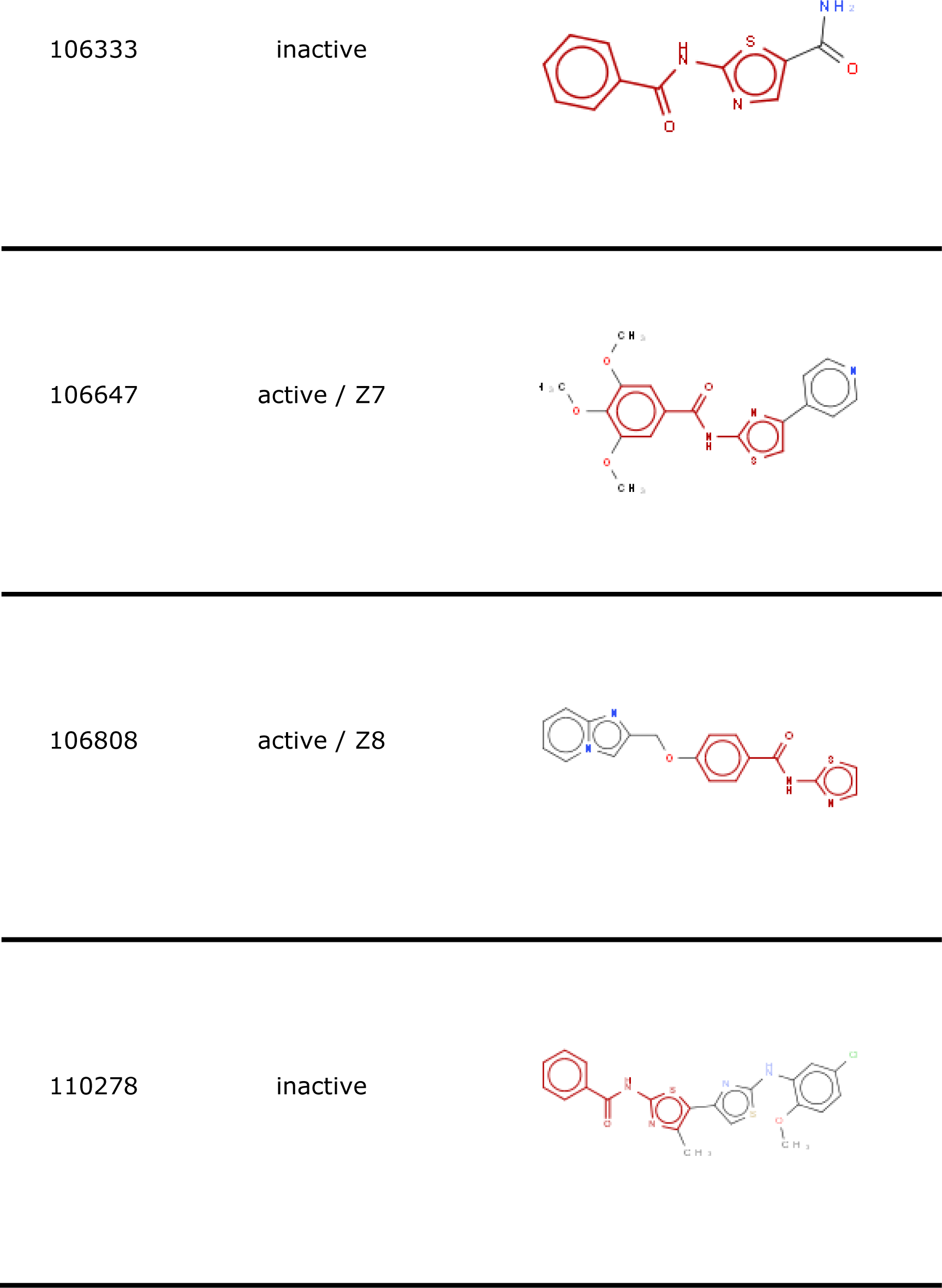
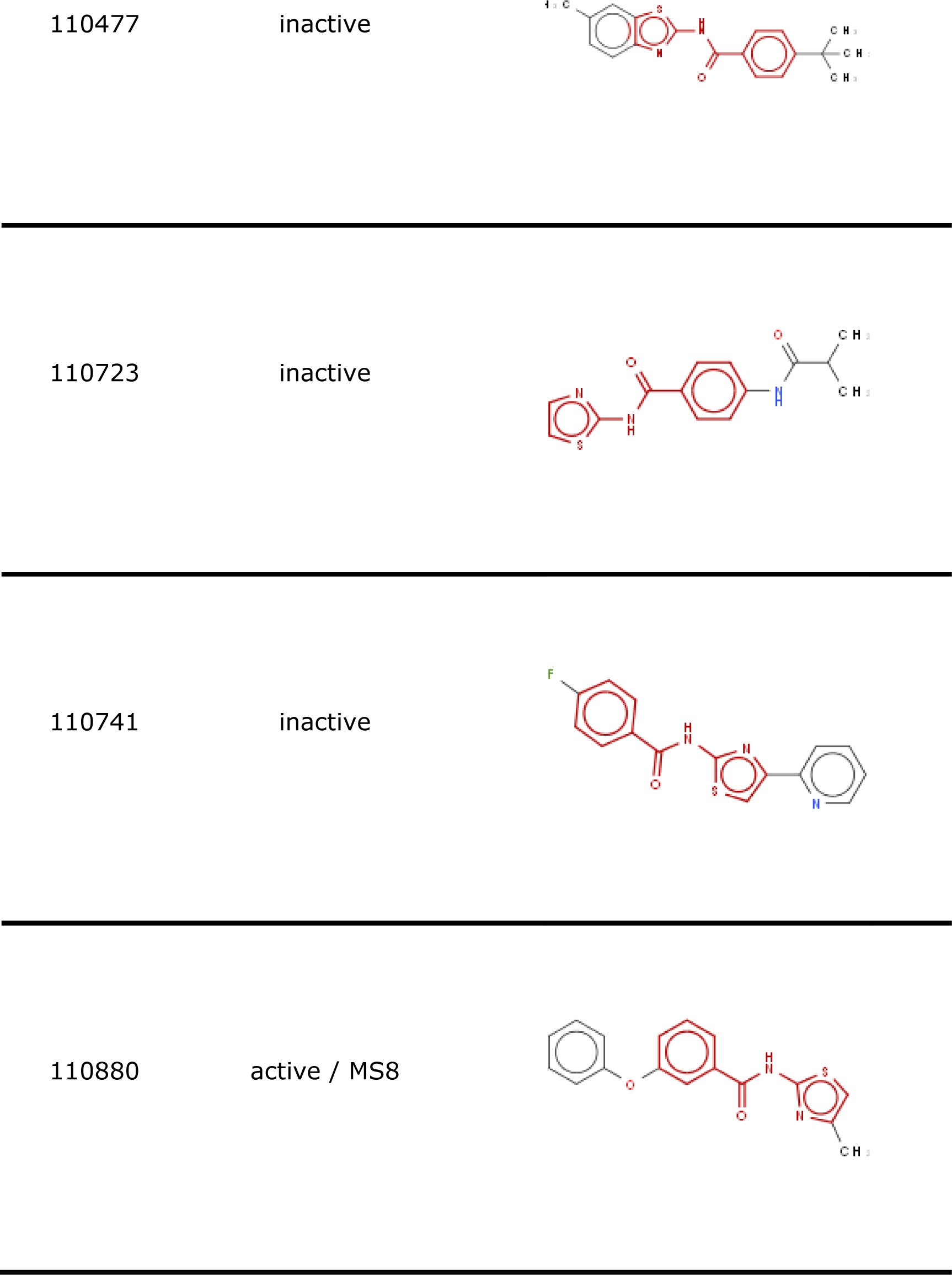
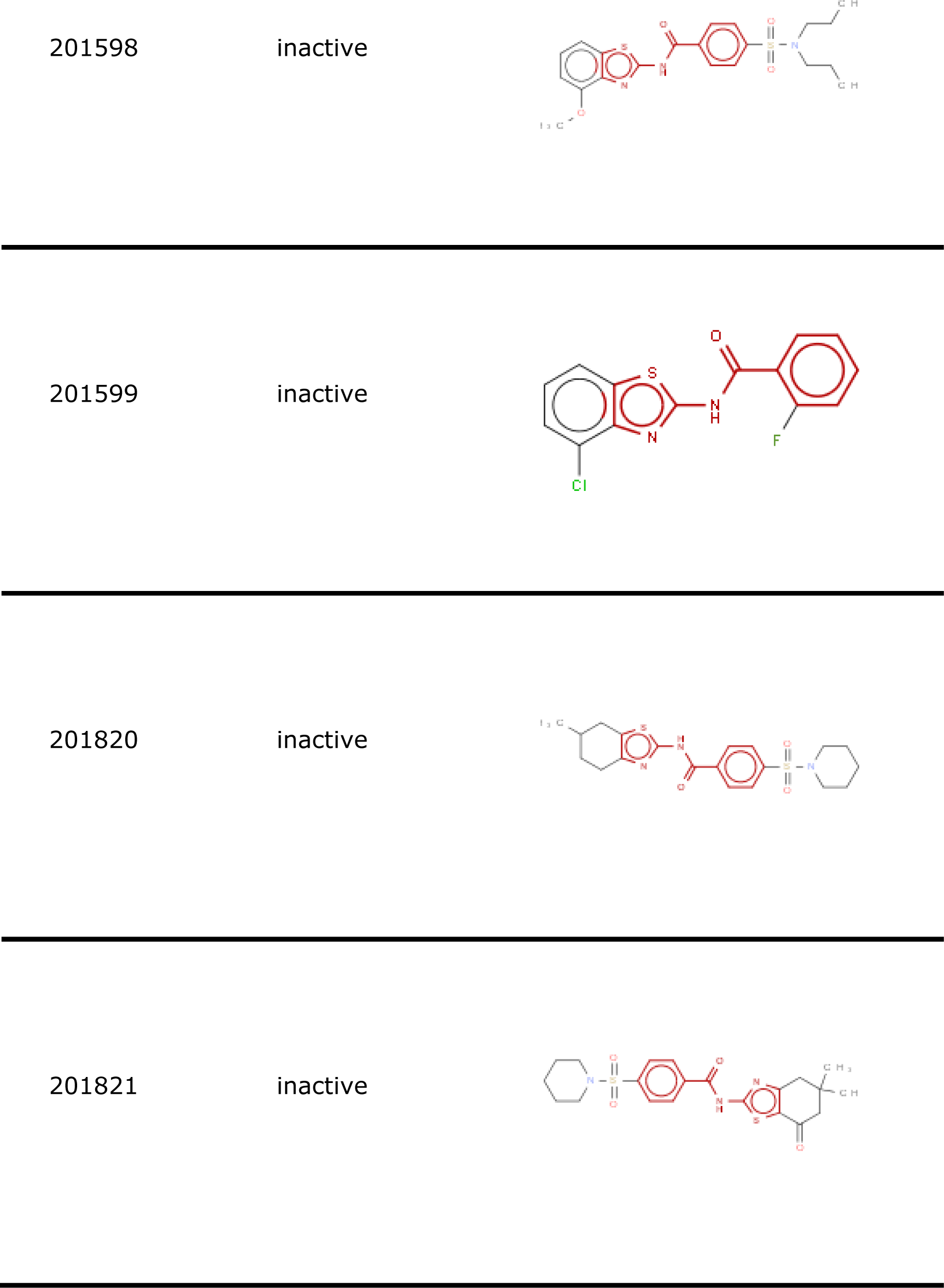
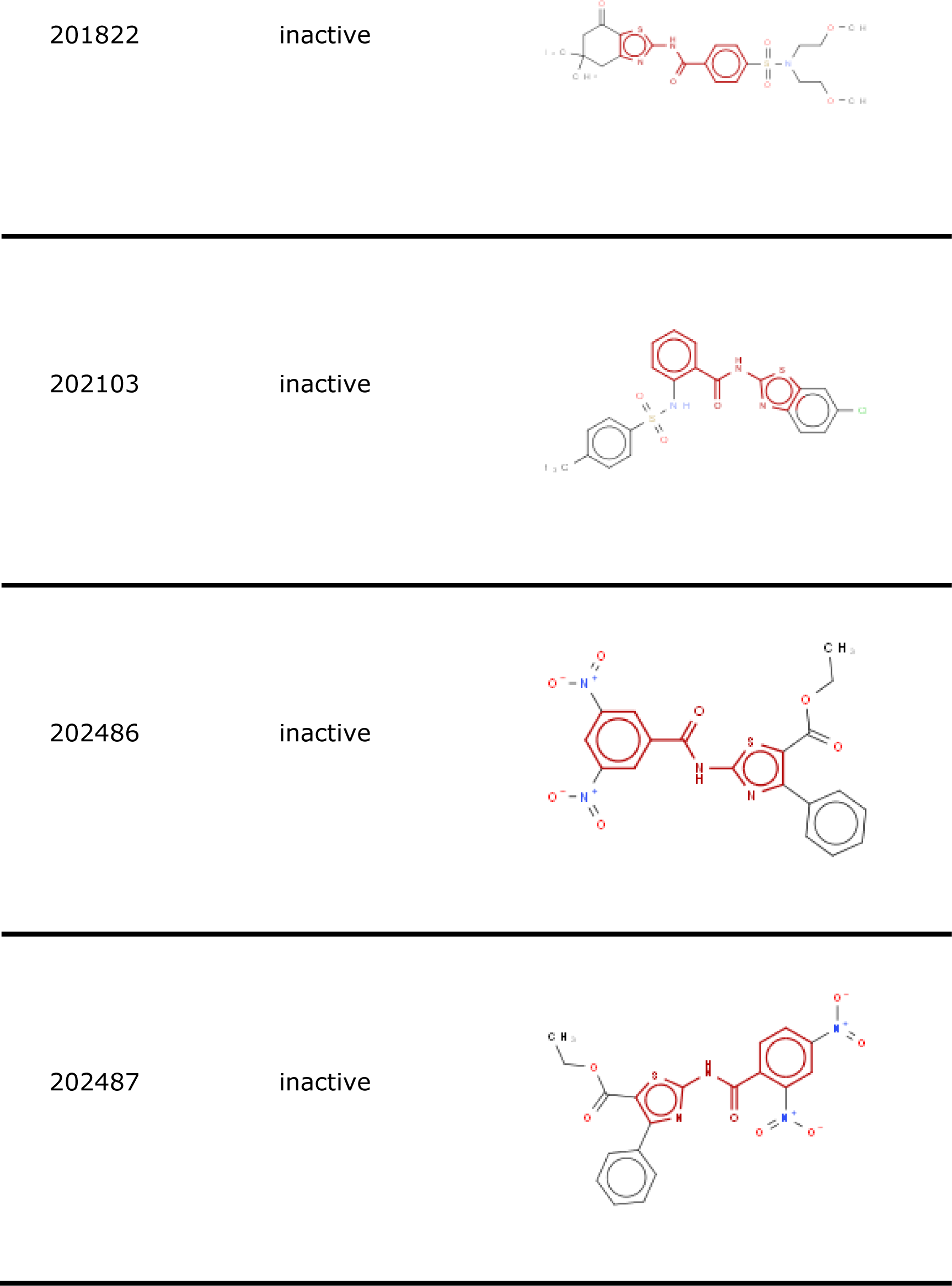
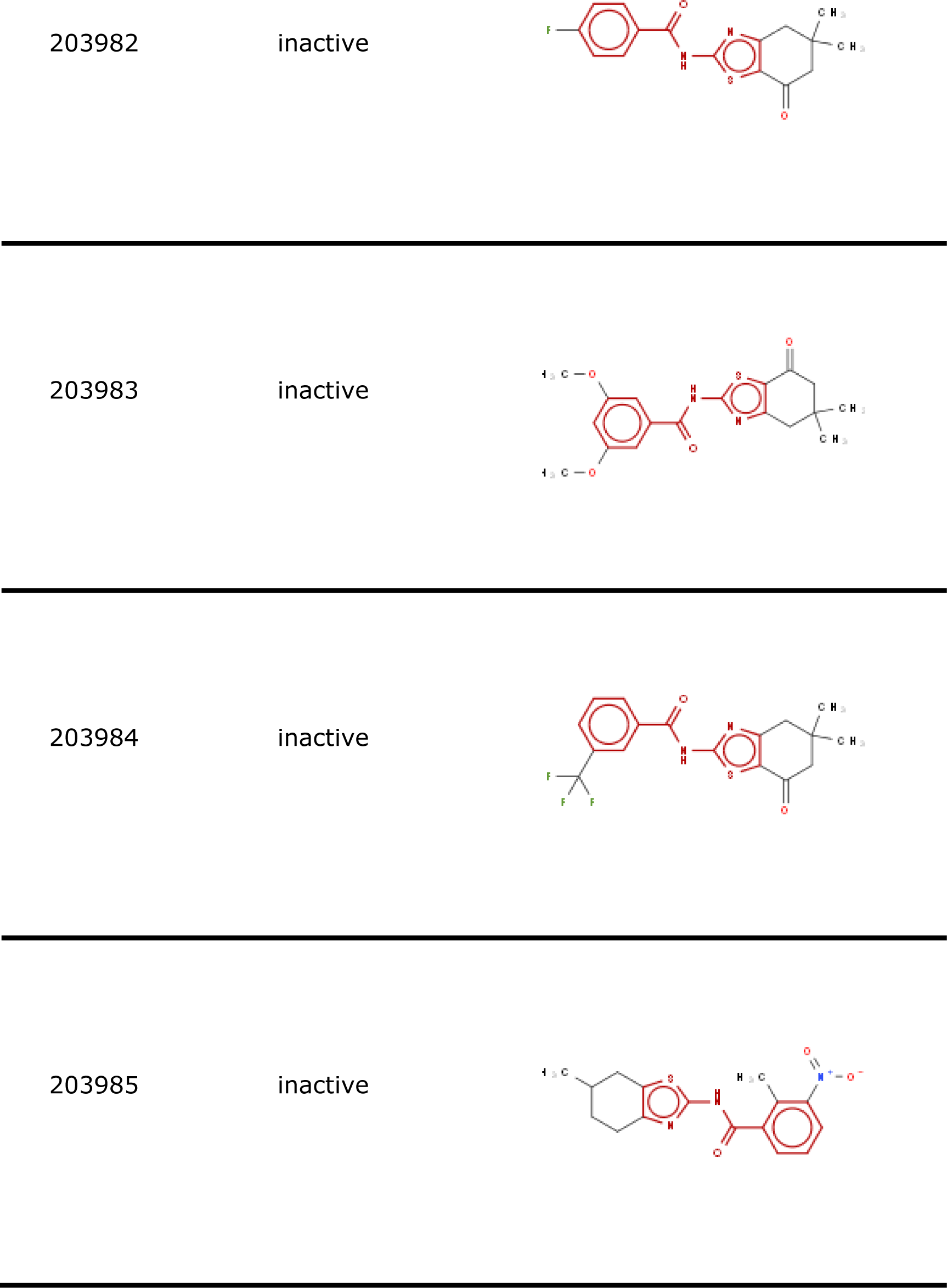
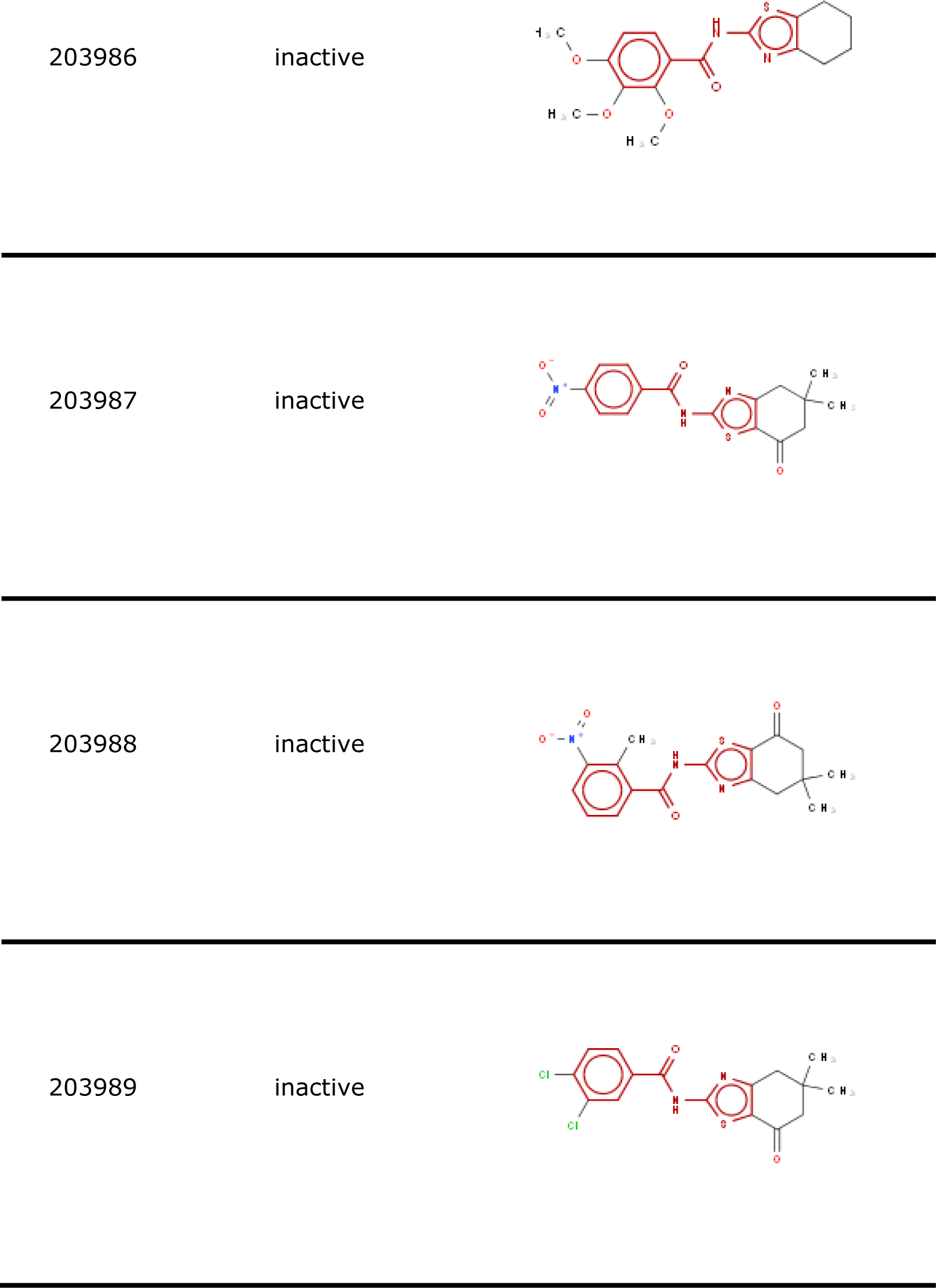
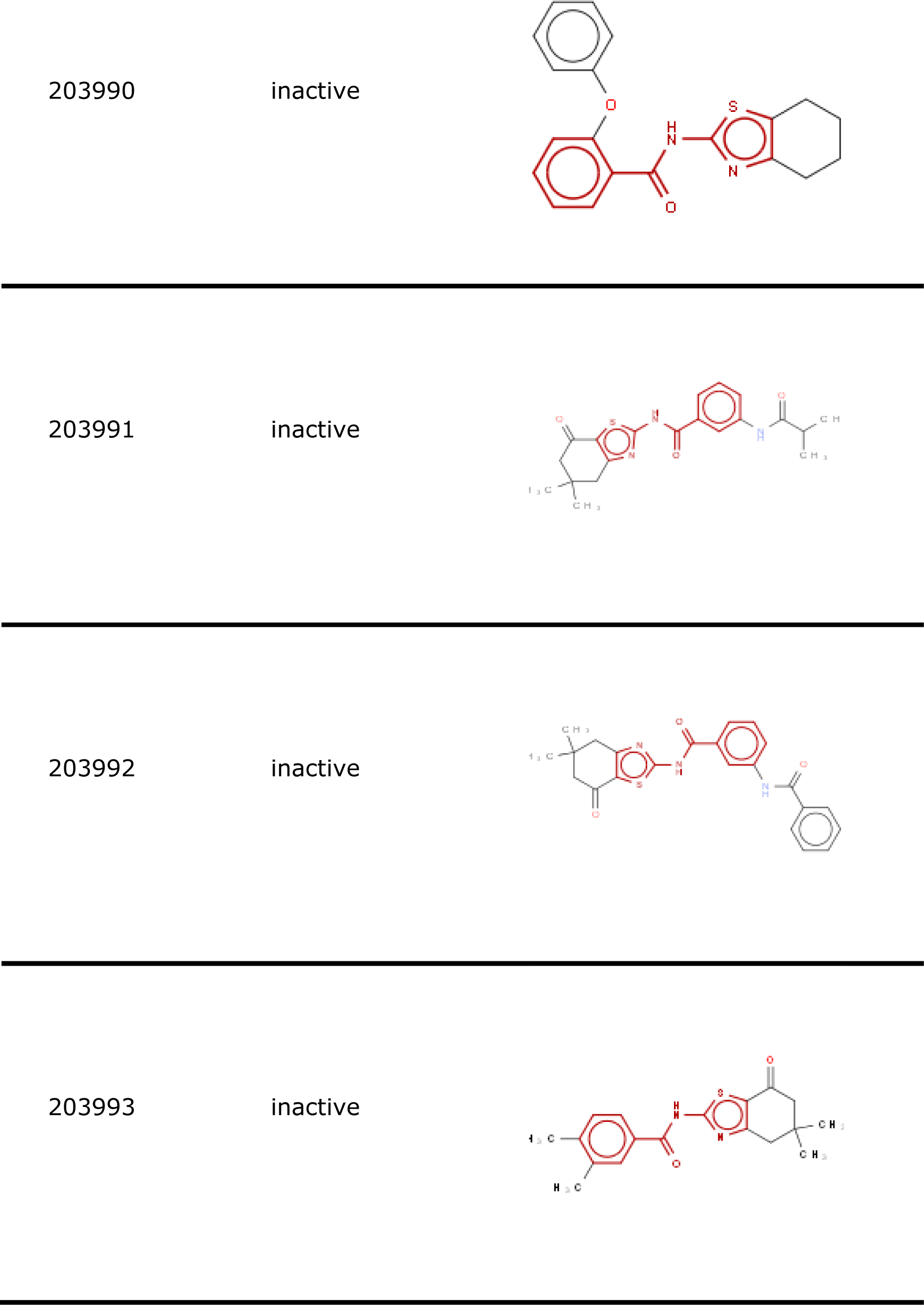
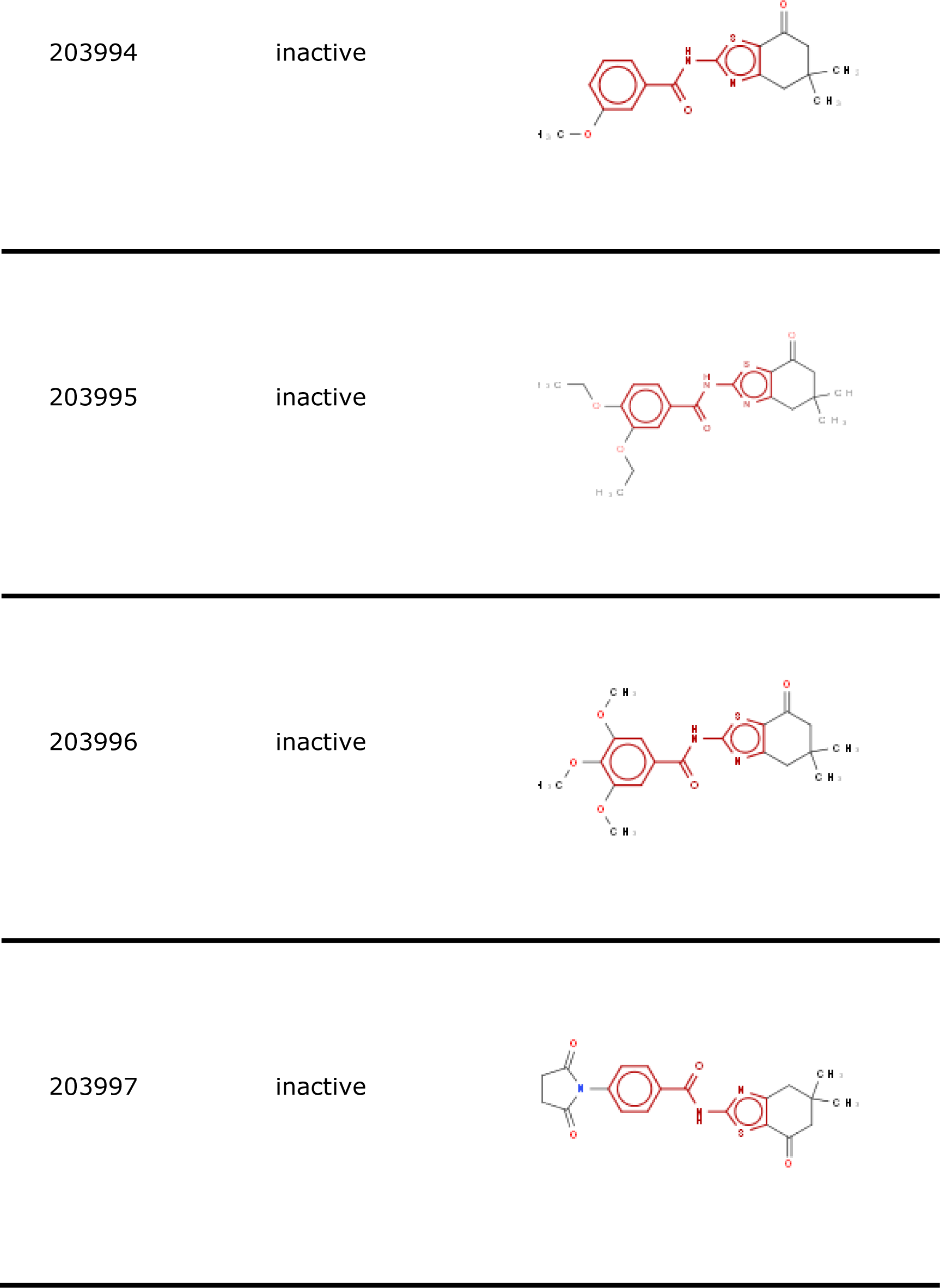
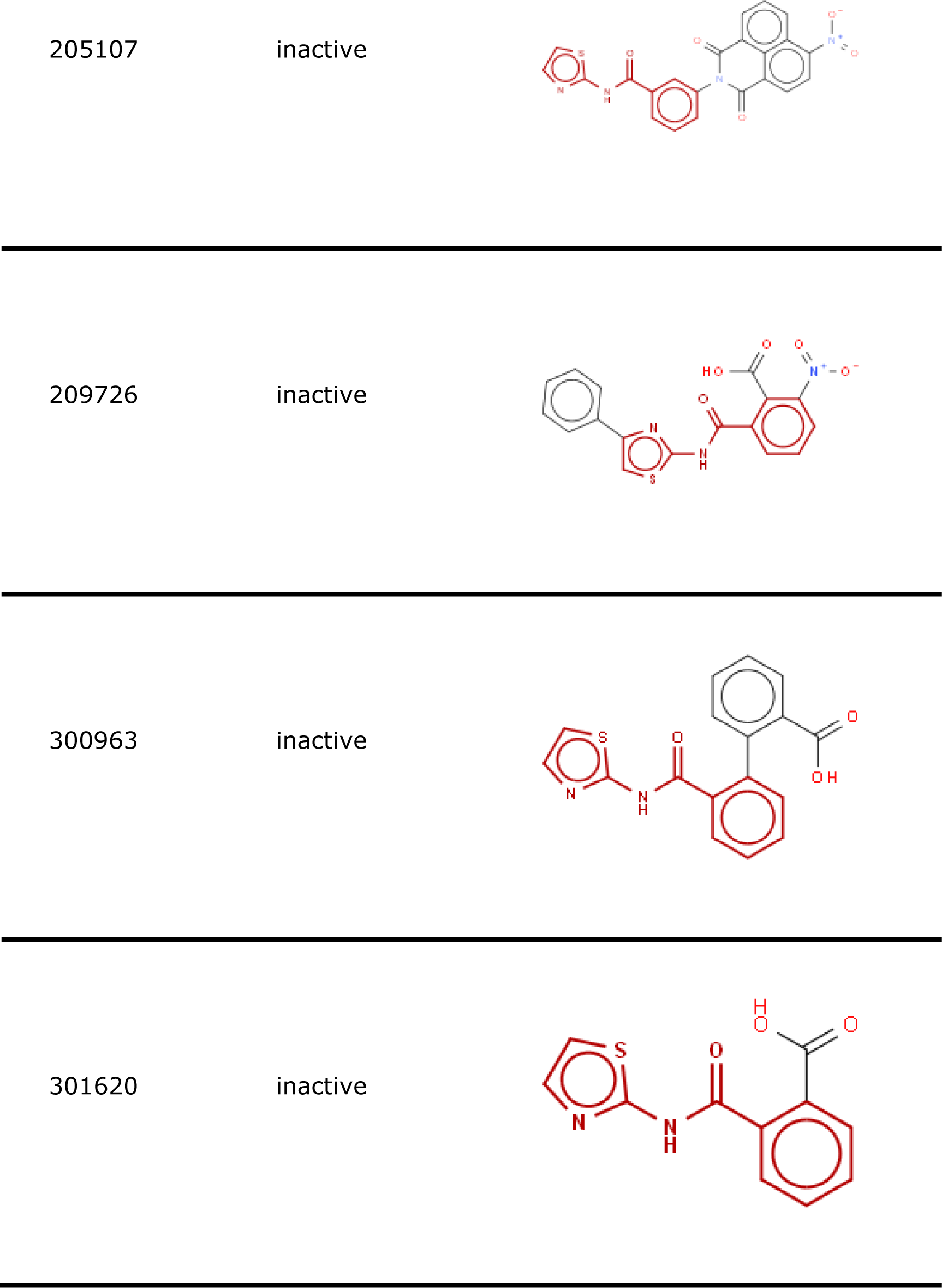
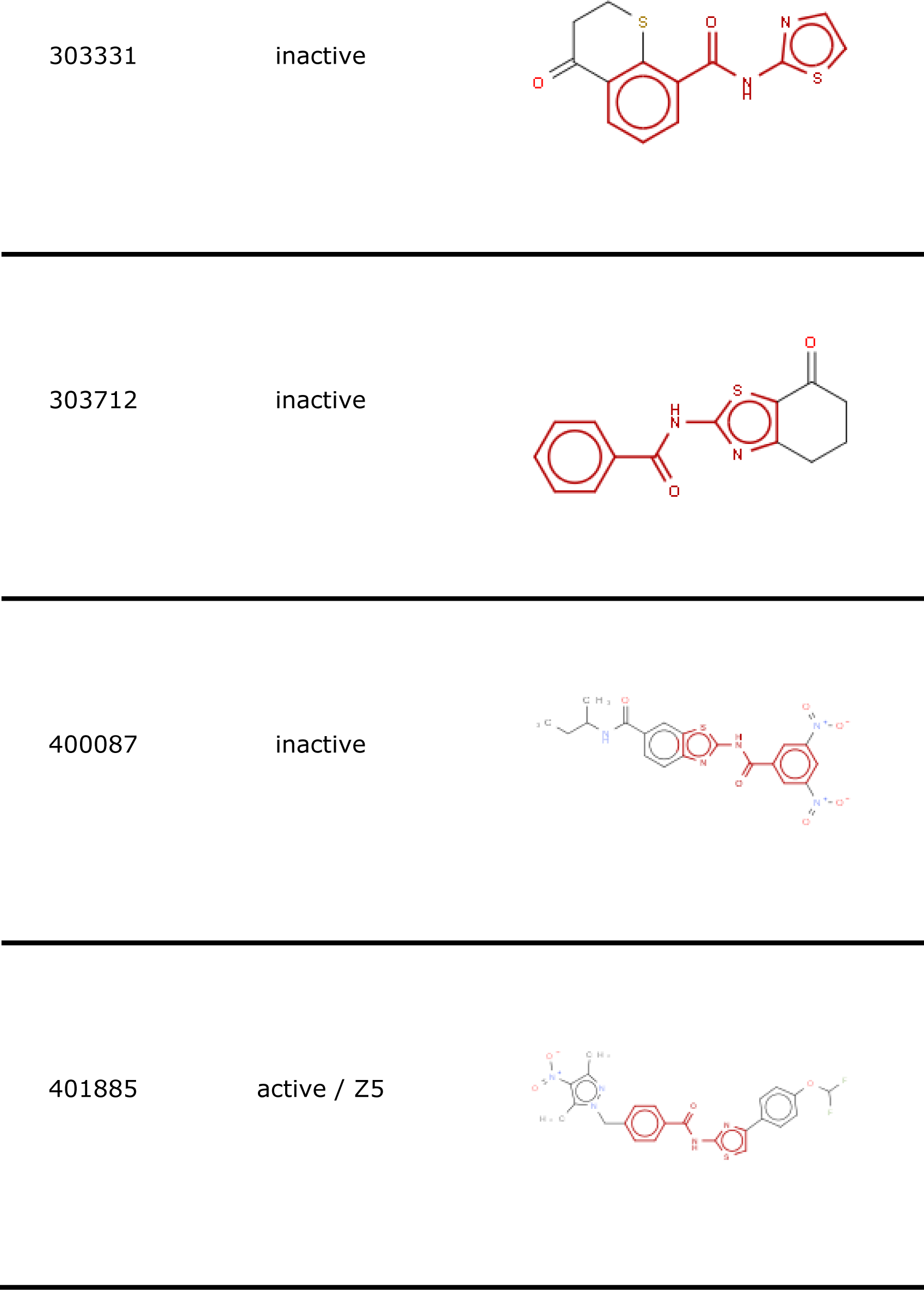
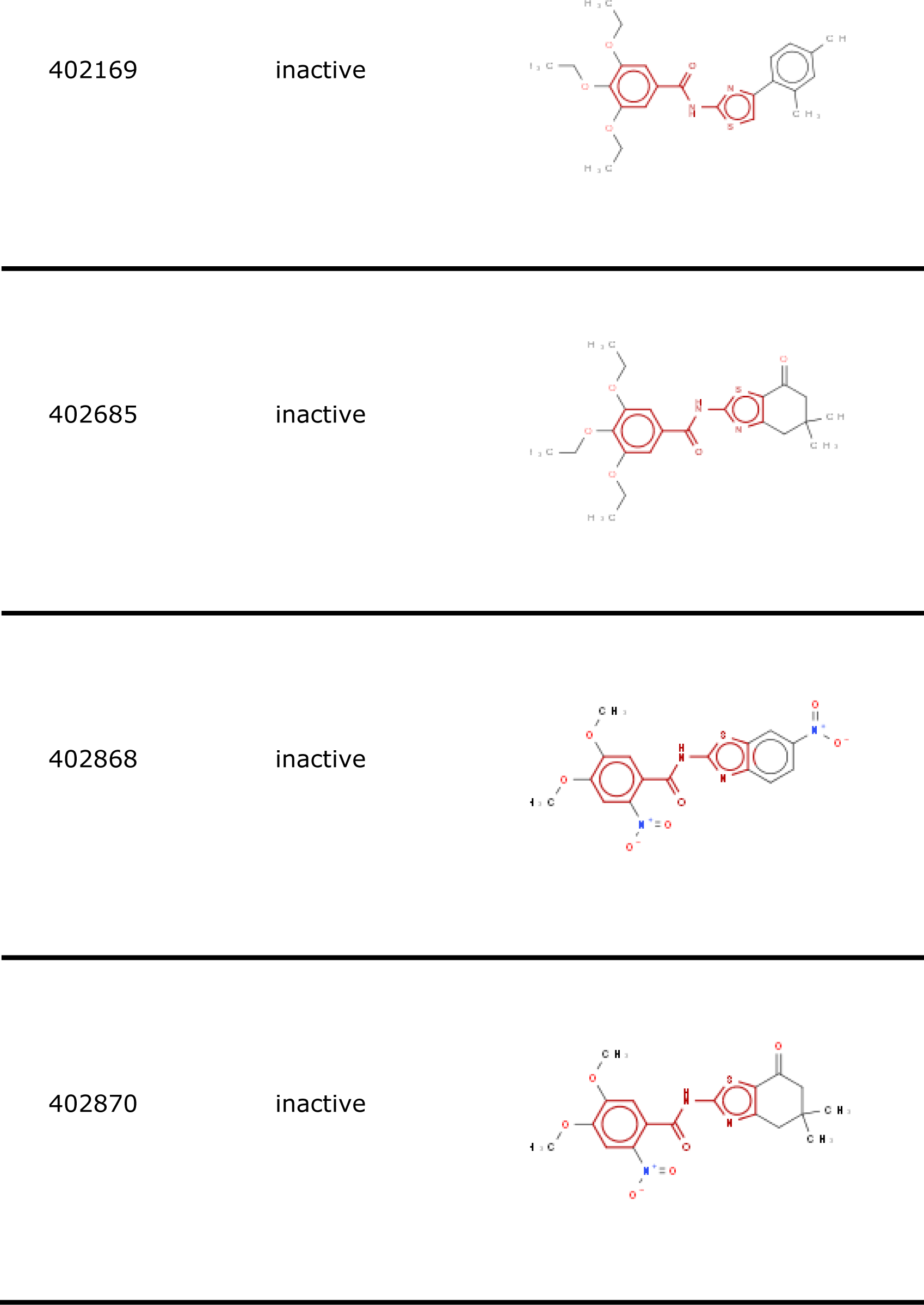
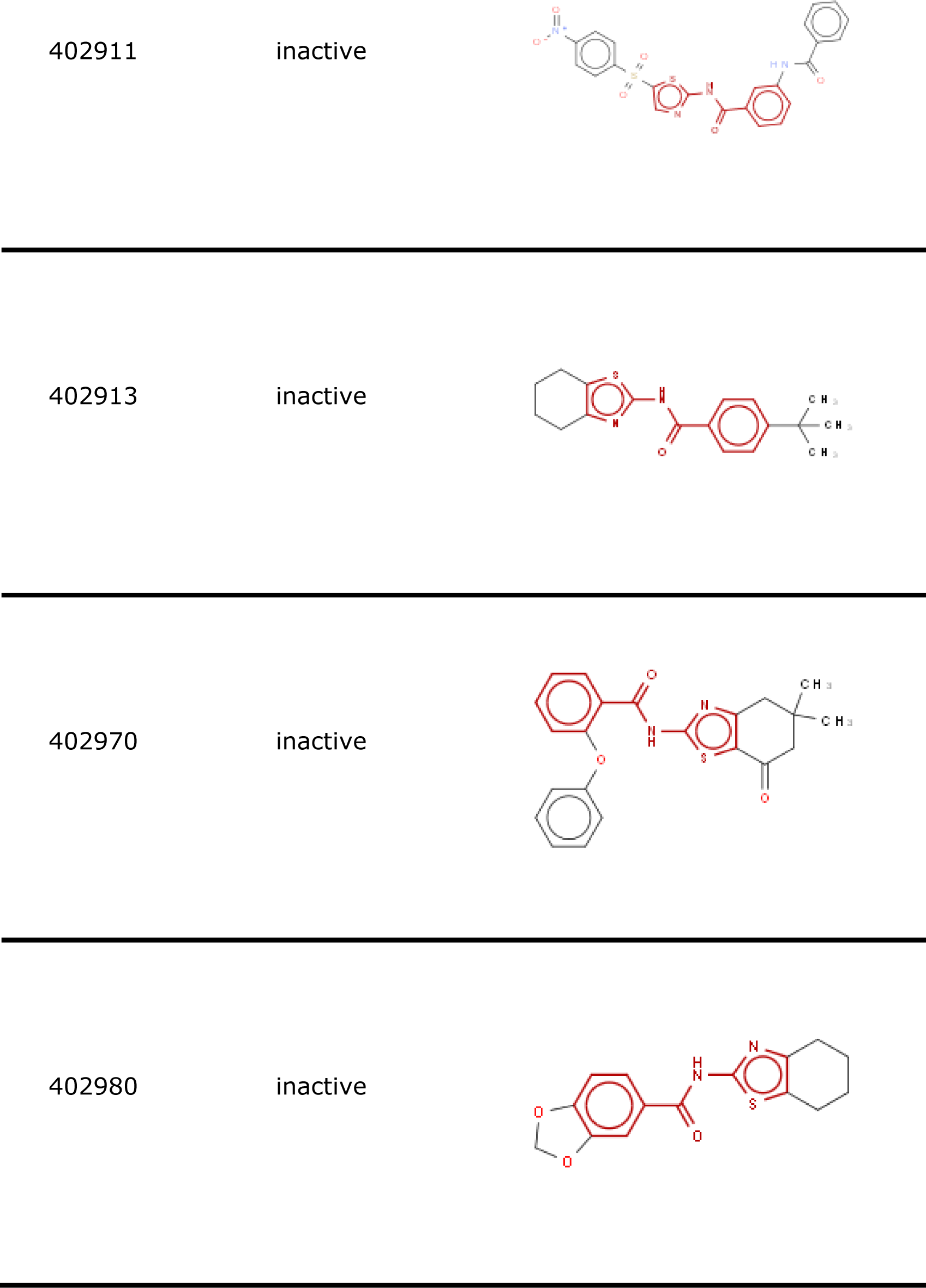
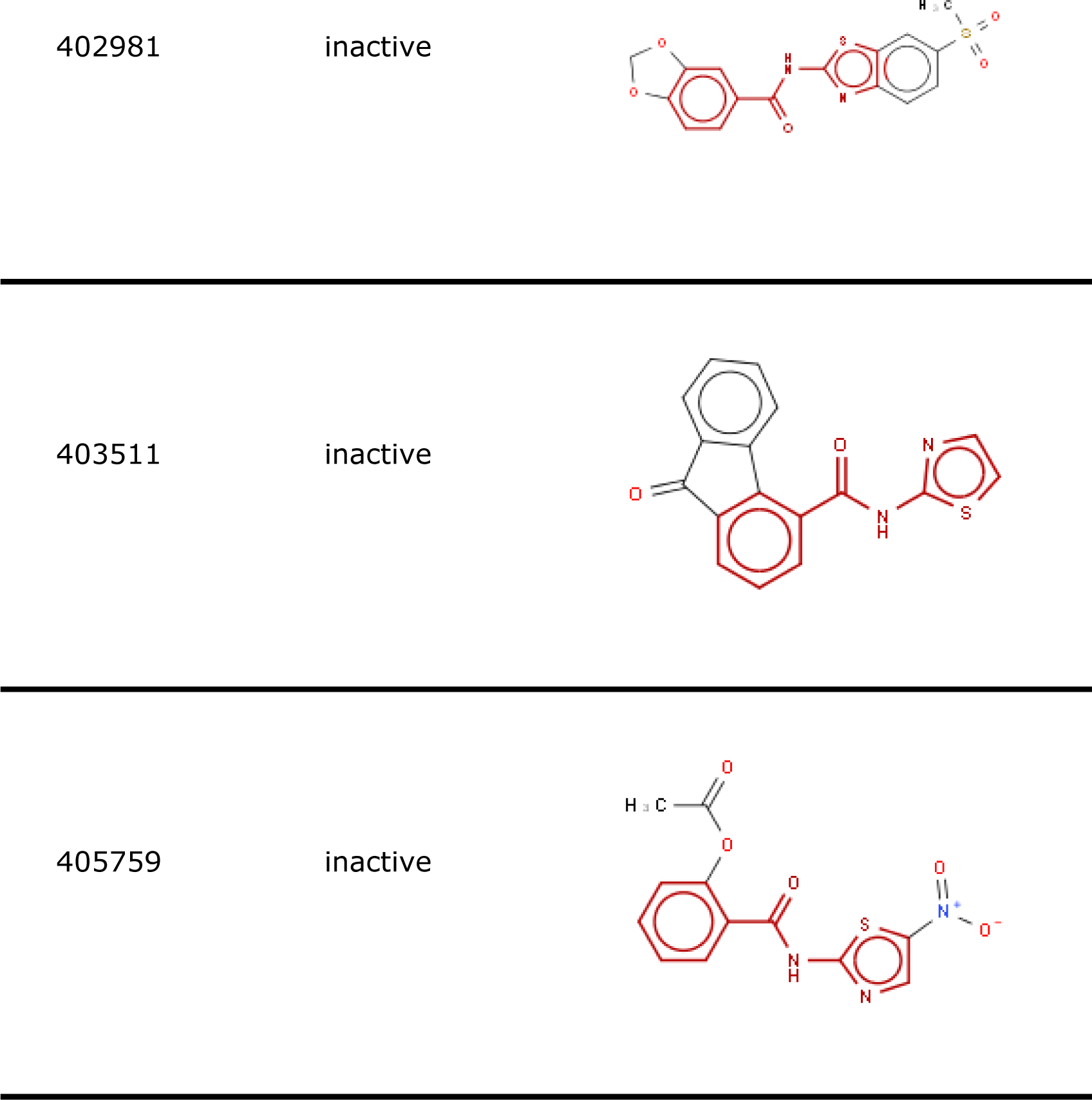
55 compounds containing the MS8 substructure highlighted in red.

